# Microbiota-derived aspartate drives pathogenic Enterobacteriaceae expansion in the inflamed gut

**DOI:** 10.1101/2022.02.14.480453

**Authors:** Woongjae Yoo, Jacob K. Zieba, Nicolas G. Shealy, Teresa P. Torres, Julia D. Thomas, Catherine D. Shelton, Nora J. Foegeding, Erin E. Olsan, Mariana X. Byndloss

## Abstract

Inflammation boosts the availability of electron acceptors in the intestinal lumen creating a favorable niche for pathogenic Enterobacteriaceae. However, the mechanisms linking intestinal inflammation-mediated changes in luminal metabolites and pathogen expansion remain unclear. Here, we show that mucosal inflammation induced by *Salmonella enterica* serovar Typhimurium (*S.* Tm) infection and chemical colitis results in increased intestinal levels of the amino acid aspartate. The *S.* Tm and *E. coli* genomes encode an aspartate ammonia-lyase (*aspA*) which converts aspartate into fumarate, an alternative electron acceptor. *S.* Tm and pathogenic *E. coli* used *aspA-*dependent fumarate respiration for growth in the murine gut only during inflammation. Such growth advantage was abolished in the gut of germ-free mice. However, mono-association of gnotobiotic mice with members of the classes Bacteroidia and Clostridia restored the benefit of aspartate utilization to the pathogens. Our findings demonstrate the role of microbiota-derived amino acids in driving respiration-dependent Enterobacteriaceae expansion during colitis.

## INTRODUCTION

The human gastrointestinal tract is colonized by a complex microbial community known as the gut microbiota. This community provides benefits to the host by contributing to nutrition and immune education (de Vos et al., 2022). In addition, the gut microbiota protects the host by limiting enteric pathogen expansion (colonization resistance) partially via nutrient sequestration (Caballero-Flores et al., 2020; Krautkramer et al., 2021; Oliphant and Allen-Vercoe, 2019). Enteric pathogens, such as *Salmonella enterica* serovar Typhimurium (*S.* Tm) and carcinogenic *pks*+ *Escherichia coli* (*E. coli*) have evolved to overcome colonization resistance by inducing and/or thriving during intestinal inflammation (colitis) (Arthur and Jobin, 2013; Arthur et al., 2014; Bronner et al., 2018; Faber et al., 2017; Shelton et al., 2022; Thiennimitr et al., 2011; Zhu et al., 2019). One of the critical mechanisms driving a luminal expansion of pathogenic Enterobacteriaceae during colitis is the generation of respiratory electron acceptors as a byproduct of the inflammatory host response. Electron acceptors enable *S*. Tm and pathogenic *E. coli* to use aerobic and anaerobic respiration to outcompete the resident microbiota and thus play a key role in altering intestinal microbiota composition during disease (Ali et al., 2014; Lopez et al., 2015, 2012; Rivera-Chavez et al., 2013; Thiennimitr et al., 2011; Winter et al., 2010). It is possible that not only the host, but also the gut microbiota may provide novel electron acceptors that support growth of enteric pathogens during gastroenteritis (Winter et al., 2010).

The crosstalk between the microbiota and enteric pathogens can be investigated by studying the flux of metabolites between microbes in the inflamed gut (Schrimpe-Rutledge et al., 2016; Tang et al., 2019; Zierer et al., 2018). Indeed, inflammation causes a significant change in the metabolic landscape of the gut, leading to the accumulation of a previously unknown set of nutrients that microorganisms inhabiting the intestinal lumen will need to compete for (Yoo et al., 2020). One metabolic interaction of interest is the microbiota’s regulation of amino acids availability and utilization. Published studies demonstrate that commensal bacteria can reduce amino acid availability in the gut lumen, limiting early intestinal colonization by enteric pathogens (Caballero-Flores et al., 2020; Nguyen et al., 2020; Popp et al., 2015; Shealy et al., 2021). Amino acids are the building blocks of proteins and serve as intermediates in energy-producing cycles like the tricarboxylic acid cycle (TCA) and the urea cycle (Akram, 2014; Morris, 2002; Salway, 2018). By sequestering amino acids, the microbiota limits protein production and potential energy production by enteric pathogens. Thus, for a pathogen to succeed in the gastrointestinal tract it must disrupt the microbiota restrictive processes such as amino acid sequestration. The mechanisms used by pathogenic Enterobacteriaceae to acquire and metabolize amino acids to fuel expansion in the inflamed gut are largely unknown.

The *S.* Tm and *E. coli* genomes encode an aspartate-ammonia lyase (*aspA*) which enables the conversion of the amino acid aspartate into fumarate, an alternative electron acceptor (Falzone et al., 1988). *S.* Tm predominantly uses the fumarate reductase encoded by *frdABCD* genes under anaerobic or hypoxic conditions for expansion through fumarate respiration in the host gut during early infection (Arguello et al., 2010; Jones et al., 2007; Mercado-Lubo et al., 2008; Paiva et al., 2009). However, the major source of fumarate during later stages of *S.* Tm intestinal infection, and whether aspartate-dependent fumarate respiration supports microbiota outgrowth by pathogenic Enterobacteriaceae in the inflamed gut remains unclear.

In this study, we show intestinal inflammation increases the abundance of the amino acid aspartate, which is dependent upon the presence of specific intestinal microbiota members. Previous studies suggest that dietary aspartate is the primary source of this amino acid in the intestinal lumen. Here, we use germ-free mouse models and dietary manipulations studies to demonstrate host and dietary aspartate do not contribute to elevated aspartate levels in the gut during inflammation. We show that *S*. Tm takes advantage of the increased aspartate pool in the intestinal lumen to outcompete the resident microbiota by converting aspartate into fumarate, an electron acceptor used for anaerobic respiration. Further, we demonstrate that increased microbiota-derived aspartate leads to carcinogenic *pks*+ *E. coli* expansion in the gut during colitis (DSS model of colitis). Lastly, our results using bacterial mutants that cannot induce intestinal inflammation suggest that aspartate-dependent fumarate respiration does not contribute to early (and inflammation-independent) intestinal colonization by pathogenic Enterobacteriaceae.

## RESULTS

### Pathogenic Enterobacteriaceae relies on aspartate for growth during colitis

Previous studies have suggested that intestinal inflammation changes the availability of amino acids in the intestinal lumen (Kitamoto et al., 2020). However, it is unclear whether infectious or noninfectious colitis may affect intestinal levels of amino acids (e.g., aspartate) used by pathogenic Enterobacteriaceae. To test the hypothesis that intestinal inflammation increases the availability of aspartate in the gut lumen, we measured aspartate levels in fecal contents of mouse models of infectious and noninfectious colitis. We used CBA/J mice infected with *S*. Tm as the model of infectious colitis, as this mouse strain develops pathogen-induced gastroenteritis in the absence of streptomycin-treatment (Faber et al., 2017; Plant and Glynn, 1976; Rivera-Chávez et al., 2016). To induce colitis independently of a pathogen (noninfectious colitis), we treated C57Bl/6J mice with 2.5% dextran sodium sulfate (DSS) in the drinking water for seven days. Indeed, we observed increased aspartate levels in fecal samples from *S*. Tm-infected mice (**Fig. 1A**) and DSS-treated mice (**Fig. 1B**), providing the initial evidence that intestinal inflammation increases aspartate availability through unknown mechanisms.

**Figure 1.**
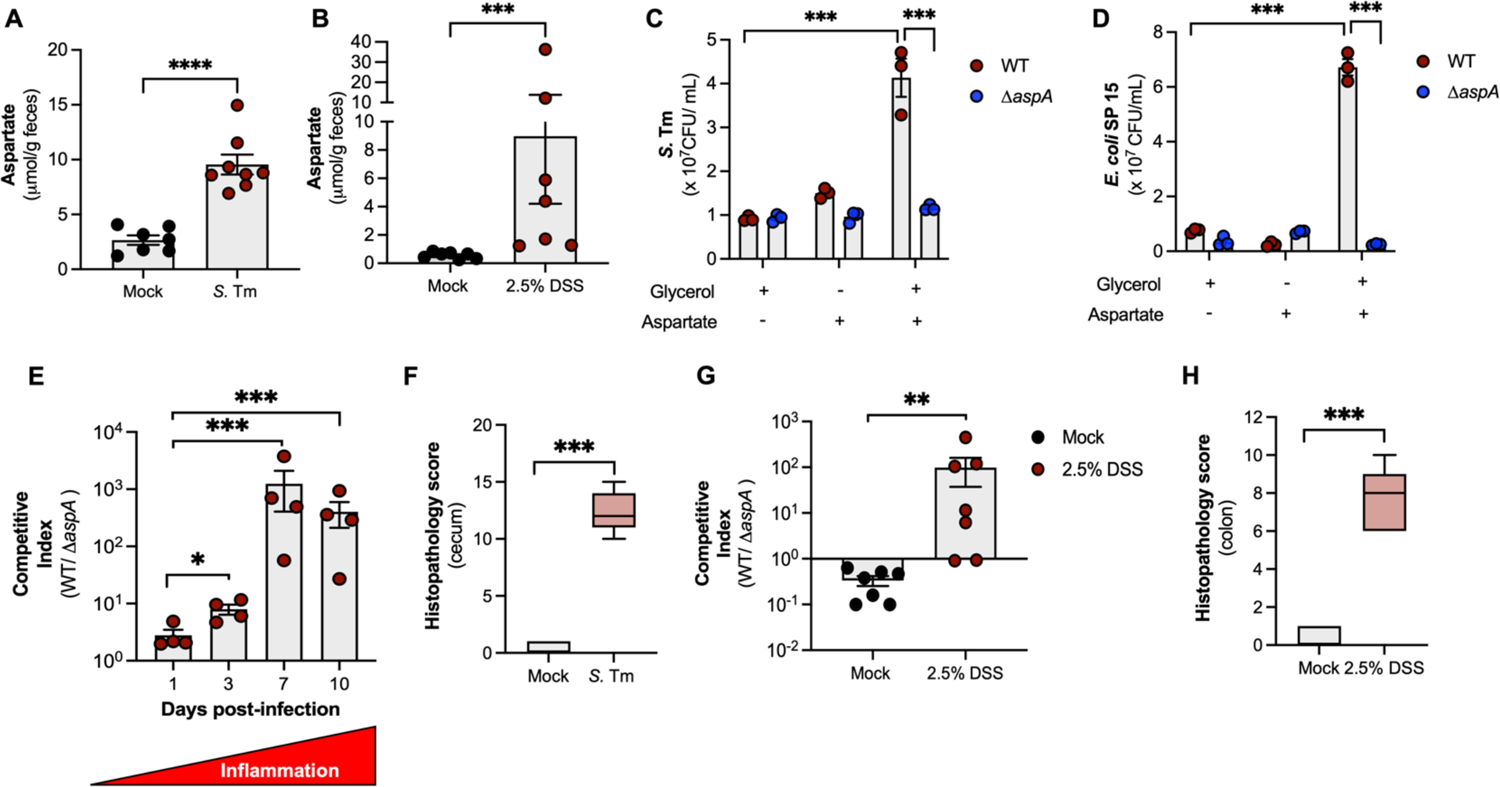
Aspartate utilization via *aspA* fuels pathogenic Enterobacteriaceae expansion during colitis. (A) Female CBA/J mice were infected intragastrically with 10^9^ CFU of *Salmonella enterica* serovar Typhimurium (*S*. Tm) for ten days. Fecal aspartate levels were measured at day ten post-infection. (B) Female C57BL/6J mice were treated with 2.5% Dextran Sodium Sulfate (DSS) in their drinking water for seven days. Fecal aspartate levels were determined seven days after beginning DSS. (C-D) Wild-type *S.* Tm (C) and *Escherichia coli* SP15 (D) and isogenic Δ*aspA* mutants were grown under anaerobic conditions in NCE minimal media supplemented or not with glycerol (40 mM) and L-aspartate (30 mM). Bacterial numbers were determined at 18 hours post-inoculation. (E) Female CBA/J mice were intragastrically infected with 10^9^ CFU of a 1:1 ratio of wild-type *S*. Tm (WT) and isogenic Δ*aspA* mutant for ten days. Competitive index in fecal samples was determined at day one, three, seven and ten post-infection. (F) Combined histopathology score of pathological lesions in the cecum of mice from (E) at ten days post-infection (n=4). (G) Female C57BL/6J mice were intragastrically infected with 10^9^ CFU of a 1:1 ratio of wild-type *E. coli* SP15 (WT) and isogenic Δ*aspA* mutant. One day after infection mice were treated with 2.5% DSS in their drinking water for seven days. Competitive index in fecal samples was determined at day eight post-infection. (H) Combined histopathology score of pathological lesions in the colon of mice from (G) at eight days post-infection (n=7). For *in vitro* experiments, each dot represents one biological replicate (average of triplicate technical replicate per biological replicate). For mouse experiments, each dot represents data from one animal (biological replicate). Bars represent mean ± SEM. The boxes in the whisker plot represent the first to third quartiles, and the mean value of the gross pathology scores is indicated by a line. **, p < 0.01; ***, p < 0.001; ****, p < 0.0001. See also Figure S1.

To investigate whether aspartate utilization conferred a fitness advantage to pathogenic Enterobacteriaceae under conditions relevant to intestinal disease (anaerobic conditions), we first generated *S*. Tm and *pks*+ *E. coli* SP15 mutants unable to use aspartate by deleting *aspA* (encoding aspartate ammonia-lyase). We determined the anaerobic growth of *S*. Tm wild-type and *ΔaspA* isogenic mutant as well as *E. coli* SP15 wild-type and Δ*aspA* isogenic mutant in minimal medium supplemented or not with 40 mM glycerol (carbon source), 30 mM aspartate and/or 40 mM nitrate (electron acceptor). Aspartate alone did not promote growth of *S.* Tm (**Fig. 1C**) or *E. coli* SP15 strains (**Fig. 1D**), nor did the addition of both nitrate and aspartate to the media (**Fig. S1A** and **S1B**). Interestingly, *S.* Tm and *E. coli* SP15 were able to grow in minimal media supplemented with glycerol and aspartate, and such phenotype was abolished in *S.* Tm and *E. coli* SP15 Δ*aspA* isogenic mutants (**Fig. 1C and 1D**). *S.* Tm and *E. coli* SP15 Δ*aspA* mutants were able to grow to similar levels as wild-type strains when glucose was given as a carbon source (**Fig. S1C and S1D**), suggesting that their inability to grow under glycerol and aspartate conditions was not due to an overall growth-defect. Taken together, our results show that pathogenic Enterobacteriaceae do not use aspartate as a carbon source for fermentation or anaerobic respiration. Instead, our results show that *S*. Tm and carcinogenic *pks*+ *E. coli* use aspartate via *aspA* to metabolize carbon sources (e.g., glycerol) via anaerobic respiration which supports the growth of pathogenic Enterobacteriaceae *in vitro* under anaerobic conditions.

The finding that the amino acid aspartate can support anaerobic-respiration dependent growth of *S.* Tm and *E. coli* SP15 *in vitro* prompted us to test if pathogenic Enterobacteriaceae can use *aspA* to take advantage of the increased levels of aspartate in the inflamed gut (**Fig. 1A-B**). First, we intragastrically infected CBA/J mice with a 1:1 ratio of wild-type *S.* Tm and an isogenic *ΔaspA* mutant and determined the competitive index between strains at day one, three, seven and ten post-infection. The growth advantage of the wild-type *S.* Tm over the *ΔaspA* mutant significantly increased as disease progressed, reaching higher levels at day ten post-infection (**Fig. 1E**) when intestinal inflammation was established (**Fig. 1F**). To confirm that intestinal inflammation supports *aspA*-dependent growth by pathogenic Enterobacteriaceae, we repeated the competition experiment in the noninfectious colitis model. We intragastrically infected C57Bl6/J mice with a 1:1 ratio of wild-type *E. coli* SP15 and an isogenic *ΔaspA* mutant one day after beginning DSS treatment. We determined the competitive index between *E. coli* SP15 strains in feces at day eight post-infection (**Fig. 1G**), when intestinal inflammation was established (**Fig. 1H**). Interestingly, the wild-type *E. coli* SP15 strain was able to significantly outcompete an isogenic *ΔaspA* mutant in the feces of DSS-treated mice, but not in mock treated animals (**Fig. 1G**). Taken together, our results suggest that inflammation-mediated changes to the intestinal environment may be necessary for aspartate-dependent intestinal expansion of pathogenic Enterobacteriaceae.

### Intestinal inflammation is required for aspartate dependent pathogenic Enterobacteriaceae expansion in the inflamed gut

The marked expansion of wild-type *S.* Tm and *E. coli* SP15 over their respective *aspA* mutants during late stages of infection (**Fig. 1E and 1G**) was surprising, as previous reports suggested that aspartate utilization is required for early intestinal colonization by Enterobacteriaceae, before the onset of inflammation (Nguyen et al., 2020; Schubert et al., 2021). Our noninfectious colitis model provided the initial evidence that pathogenic Enterobacteriaceae does not rely on *aspA*-dependent aspartate utilization for growth in the colonic lumen in the absence of inflammation (**Fig. 1G-H**). To further assess whether intestinal inflammation is indeed required to support aspartate-dependent expansion of pathogenic Enterobacteriaceae, we repeated our infectious colitis mouse model (CBA/J mice) using an inflammation-deficient *S*. Tm strain (Δ*invA* Δ*spiB). S.* Tm uses two type III secretion systems (T3SS) to invade the intestinal epithelium and perform intracellular replication (Galán and Curtiss, 1989). A mutant strain (Δ*invA* Δ*spiB*) of *S.* Tm is defective in both T3SS and does not cause inflammation in a mouse model (Coombes et al., 2005; Raffatellu et al., 2009). Thus, we constructed an Δ*aspA* mutant in the *S.* Tm inflammation-deficient strain and then infected CBA/J mice with an equal mixture of Δ*invA* Δ*spiB* and Δ*invA* Δ*spiB* Δ*aspA*. The competitive index was determined one, three, seven and ten days after infection. In contrast to the competitive advantage observed for wild-type *S.* Tm over Δ*aspA,* no competitive advantage was observed for Δ*invA* Δ*spiB* over Δ*invA* Δ*spiB* Δ*aspA,* revealing that inflammation is required for aspartate to be advantageous to *S.* Tm in the intestinal lumen (**Fig. 2A**). Histopathology analysis confirmed that infection with *S*. Tm Δ*invA* Δ*spiB* did not induce intestinal inflammation (**Fig. 2B**). The concentration of aspartate was measured in the feces of mice infected with wild-type or Δ*invA* Δ*spiB* and a significant difference in fecal aspartate levels was observed (**Figure 2C**) indicating that the lack of an advantage of Δ*invA* Δ*spiB* over Δ*invA* Δ*spiB* Δ*aspA* could be due to decreased levels of aspartate in Δ*invA* Δ*spiB*-infected mice. To address this concern, we repeated the infectious colitis experiment and increased intestinal concentrations of aspartate in Δ*invA* Δ*spiB*-infected mice by providing aspartate (0.4%) in the drinking water throughout the experiment (**Fig. 2D-F**). The aspartate supplementation restored the ability of the inflammation-deficient *S.* Tm strain to outcompete an isogenic *ΔaspA* mutant at ten days post-infection (**Fig. 2D**), in the absence of intestinal inflammation (**Fig. 2E**). Aspartate supplementation in the drinking water significantly increased aspartate levels in the intestinal lumen of *S.* Tm Δ*invA* Δ*spiB*-infected mice (**Fig. 2F**). Our results demonstrate that inflammation-mediated increase in aspartate levels drive expansion of pathogenic Enterobacteriaceae in the gut lumen during colitis.

**Figure 2.**
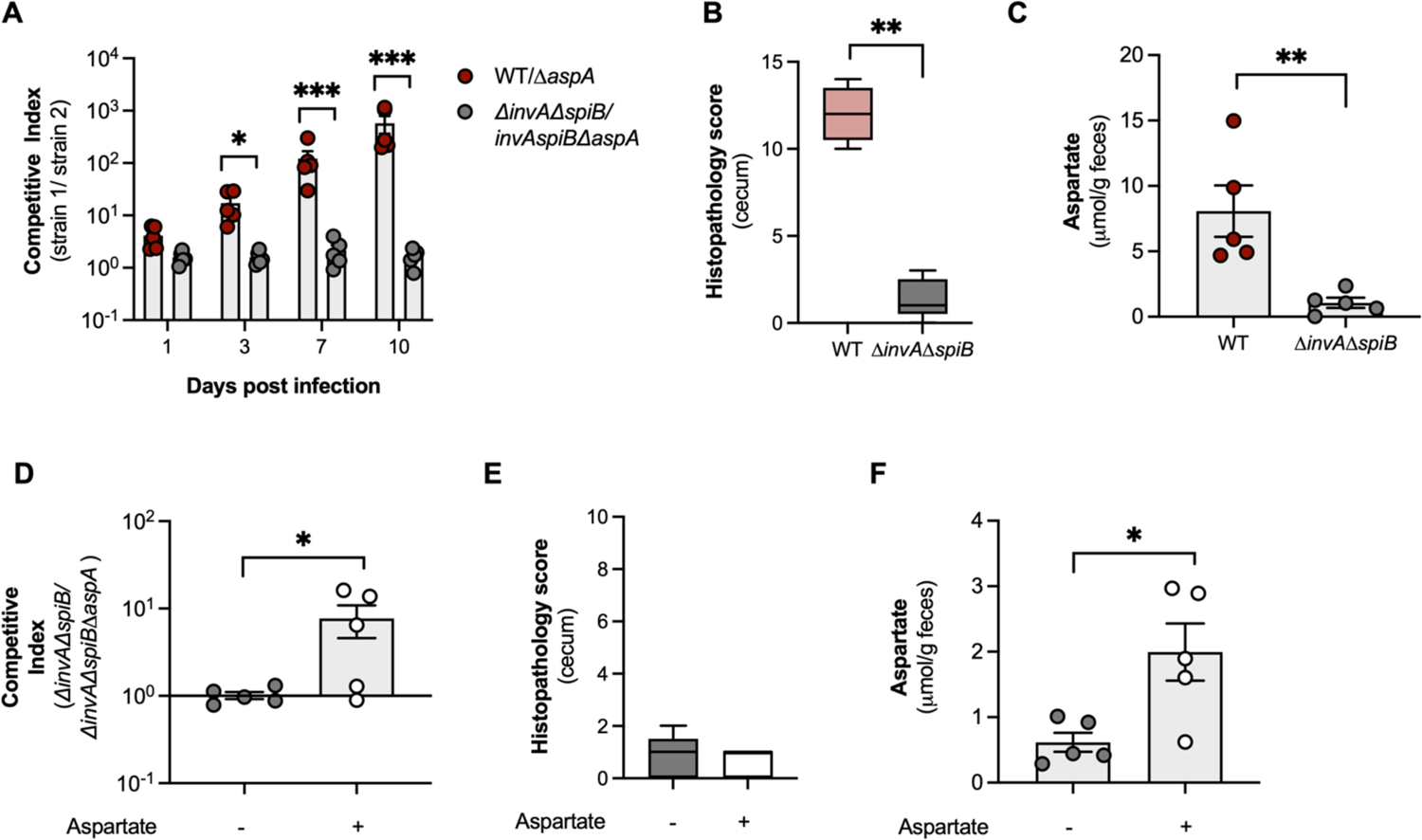
Intestinal inflammation is required for aspartate utilization by *S*. Tm in the gut. (A) Female CBA/J mice were intragastrically infected with 10^9^ CFU of a 1:1 ratio of wild-type *S*. Tm (WT) and isogenic Δ*aspA* mutant or a 1:1 ratio of *S*. Tm Δ*invA* Δ*spiB* and Δ*invA* Δ*spiB* Δ*aspA* isogenic mutant for ten days. Competitive index in fecal samples was determined at day one, three, seven and ten post-infection. (B) Combined histopathology score of pathological lesions in the cecum of mice from (B) at ten days post-infection (n=5). (C) Fecal aspartate levels from mice in (A) measured at day ten post-infection. (D) Female CBA/J mice were intragastrically infected with 10^9^ CFU of a 1:1 ratio of *S*. Tm Δ*invA* Δ*spiB* and Δ*invA* Δ*spiB* Δ*aspA* isogenic mutant for ten days. A subset of mice received L-aspartate (0.4%) supplementation in their drinking water for the duration of the experiment (Aspartate +). (E) Combined histopathology score of pathological lesions in the cecum of mice from (B) at ten days post-infection (n=5). (F) Fecal aspartate levels from mice in (A) measured at day ten post-infection. Each dot represents data from one animal (biological replicate). Bars represent mean ± SEM. The boxes in the whisker plot represent the first to third quartiles, and the mean value of the gross pathology scores is indicated by a line. *, p<0.05; **, p < 0.01; ***, p < 0.001.

### Chemotaxis towards and transport of exogenous aspartate is required for pathogenic Enterobacteriaceae expansion in the inflamed gut

The genome of pathogenic Enterobacteriaceae encodes genes that enable chemotaxis towards aspartate and uptake of this amino acid. Methyl-accepting chemotaxis protein ΙΙ (encoded by *tar*) facilitates bacterial movement towards aspartate (Slocum and Parkinson, 1985), while C4-dicarboxylate transporter (encoded by *dcuA*) uptakes exogenous aspartate and other dicarboxylates such as malate (Nguyen et al., 2020). Thus, we sought to investigate whether genes related to aspartate chemotaxis, uptake and utilization would be affected in pathogenic Enterobacteriaceae grown in the presence of aspartate under anaerobic conditions. We determined the expression of the genes *tar*, *aspA* and *ducA* of wild-type *S*. Tm grown in minimal medium supplemented or not with 40 mM glycerol (carbon source), 30 mM aspartate and/or 40 mM nitrate (electron acceptor). Growth of *S.* Tm on aspartate alone or in combination with glycerol resulted in increased expression of *tar*, *aspA* and *dcuA* when compared to *S.* Tm grown on glycerol alone (**Fig. 3A**). Additionally, growth of *S*. Tm on aspartate in the presence of glycerol resulted in increased expression of *dcuA* and *aspA* when compared to bacteria grown in aspartate alone (**Fig. 3A**), confirming that expression of aspartate uptake and utilization genes is induced under aspartate-dependent anaerobic respiration conditions *in vitro*.

**Figure 3.**
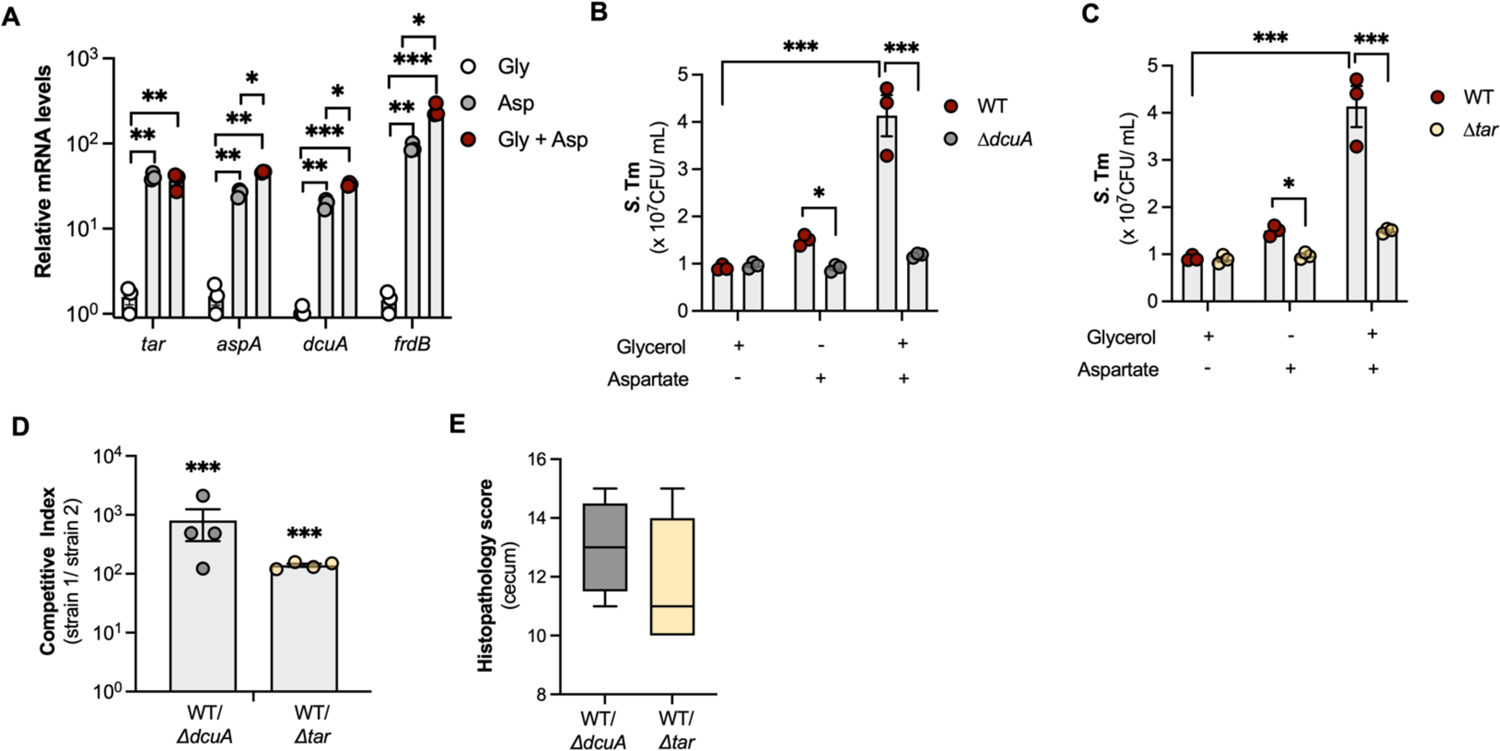
Transport of exogenous aspartate and chemotaxis towards this amino acid support *S*. Tm expansion in the inflamed gut. (A) Wild-type *S.* Tm was grown in NCE minimal media supplemented or not with glycerol (40 mM) and L-aspartate (30 mM) under anaerobic conditions. Gene expression analysis of the aspartate transport and utilization genes *tar, aspA, dcuA* and *frdB* was performed by qRT-PCR at four hours post-inoculation, and target gene transcription of each sample was normalized to 16S rRNA mRNA levels. (B-C) Wild-type *S.* Tm and isogenic *ΔdcuA* (B) and *Δtar* (C) mutants were grown in NCE minimal media supplemented or not with glycerol (40 mM) and L-aspartate (30 mM) for 18 hours under anaerobic conditions. (D) Female CBA/J mice were intragastrically infected with 10^9^ CFU of a 1:1 ratio of wild-type *S*. Tm (WT) and isogenic *ΔdcuA* mutant or a 1:1 ratio of wild-type *S*. Tm (WT) and isogenic *Δtar* mutant for ten days. Competitive index in fecal samples was determined at day ten post-infection. (E) Combined histopathology score of pathological lesions in the cecum of mice from (D) at ten days post-infection (n=4). Bars represent mean ± SEM. The boxes in the whisker plot represent the first to third quartiles, and the mean value of the gross pathology scores is indicated by a line. *, p<0.05; **, p < 0.01; ***, p < 0.001.

Next, we determined whether transport of exogenous aspartate or chemotaxis towards this amino acid conferred a fitness advantage to *S.* Tm *in vitro* under anaerobic conditions. We assessed the anaerobic growth of *S*. Tm wild-type and *ΔdcuA* and *Δtar* isogenic mutants in minimal medium supplemented or not with 40 mM glycerol (carbon source) and 30 mM aspartate. *S.* Tm was able to grow in minimal medium supplemented with glycerol and aspartate (**Fig. 3B-C**), and such phenotype was abrogated in *S.* Tm isogenic *ΔdcuA* (**Fig. 3B**) and *Δtar* (**Fig. 3C**) isogenic mutants. *S.* Tm *ΔdcuA* and *Δtar* mutants were able to grow to similar levels as wild-type strains when glucose was given as a carbon source (data not shown), suggesting that their inability to grow under glycerol and aspartate conditions was not due to an overall growth defect. Taken together, our results show that transport of exogenous aspartate and chemotaxis towards this amino acid support respiration-dependent growth of pathogenic Enterobacteriaceae *in vitro* under anaerobic conditions.

Our *in vitro* results led us to investigate whether transport of exogenous aspartate and chemotaxis towards this amino acid enable pathogenic Enterobacteriaceae to take advantage of the increased levels of aspartate in the inflamed gut (**Fig. 1A-B**). To do so, we intragastrically infected CBA/J mice with a 1:1 ratio of wild-type *S.* Tm and an isogenic *ΔdcuA* mutant or a 1:1 ratio of wild-type *S.* Tm and an isogenic *Δtar* mutant and determined the competitive index between strains at day 10 post-infection. Wild-type *S.* Tm was able to significantly outcompete the *ΔducA* mutant as well as the *Δtar* mutant (**Fig. 3D**) in fecal samples from infected mice. Such increased competitive advantage of wild-type *S*. Tm over the *ΔdcuA* and *Δtar* mutants was associated with the presence of marked intestinal inflammation (**Fig. 3E**), suggesting that the chemotaxis towards and transport of exogenous aspartate promotes the expansion of pathogenic Enterobacteriaceae during infectious colitis.

### Aspartate supports intestinal expansion of enteric pathogens by enabling anaerobic fumarate respiration

Pathogenic Enterobacteriaceae have evolved to use fumarate reductase, encoded by *frdABCD* genes, to grow under anaerobic conditions *in vitro* and *in vivo* (Iverson et al., 1999; Jones et al., 2007; Mercado-Lubo et al., 2008; Paiva et al., 2009). Interestingly, the *S.* Tm and *pks*+ *E. coli* SP15 genomes encode the *aspA* gene (aspartate ammonia-lyase), which enable the conversion of the amino acid aspartate into fumarate (Nguyen et al., 2020; Nógrády et al., 2003) (**Fig. S2A**). Indeed, anaerobic growth of *S*. Tm in the presence of aspartate alone or in combination with glycerol induced expression of one of the genes in the fumarate reductase operon, *frdB* (**Fig. 3A**). To investigate whether aspartate utilization conferred a fitness advantage to pathogenic Enterobacteriaceae by supporting fumarate respiration, we determined the anaerobic growth of *S*. Tm wild-type and *ΔfrdABCD* (*Δfrd*) isogenic mutants as well as *E. coli* SP15 wild-type and *Δfrd* isogenic mutants in minimal medium supplemented or not with 40 mM glycerol (carbon source) and 30 mM aspartate (**Fig. 4A-B**). Aspartate alone did not promote growth of *S.* Tm (**Fig. 4A**) *or E. coli* SP15 strains (**Fig. 4B**) in minimal medium. However, *S.* Tm and *E. coli* SP15 were able to grow in minimal medium supplemented with glycerol and aspartate, and such phenotype was abolished in *S.* Tm and *E. coli* Δ*frd* isogenic mutants (**Fig. 4A and 4B**). *S.* Tm and *E. coli* Δ*frd* mutants were able to grow to similar levels as wild-type strains when glucose was given as a carbon source (data not shown), suggesting that their inability to grow under glycerol and aspartate conditions was not due to an overall growth defect. These results confirm previous reports (Nguyen et al., 2020; Schubert et al., 2021) that pathogenic Enterobacteriaceae utilize aspartate as a source of fumarate to support respiration-dependent growth under *in vitro* anaerobic conditions.

**Figure 4.**
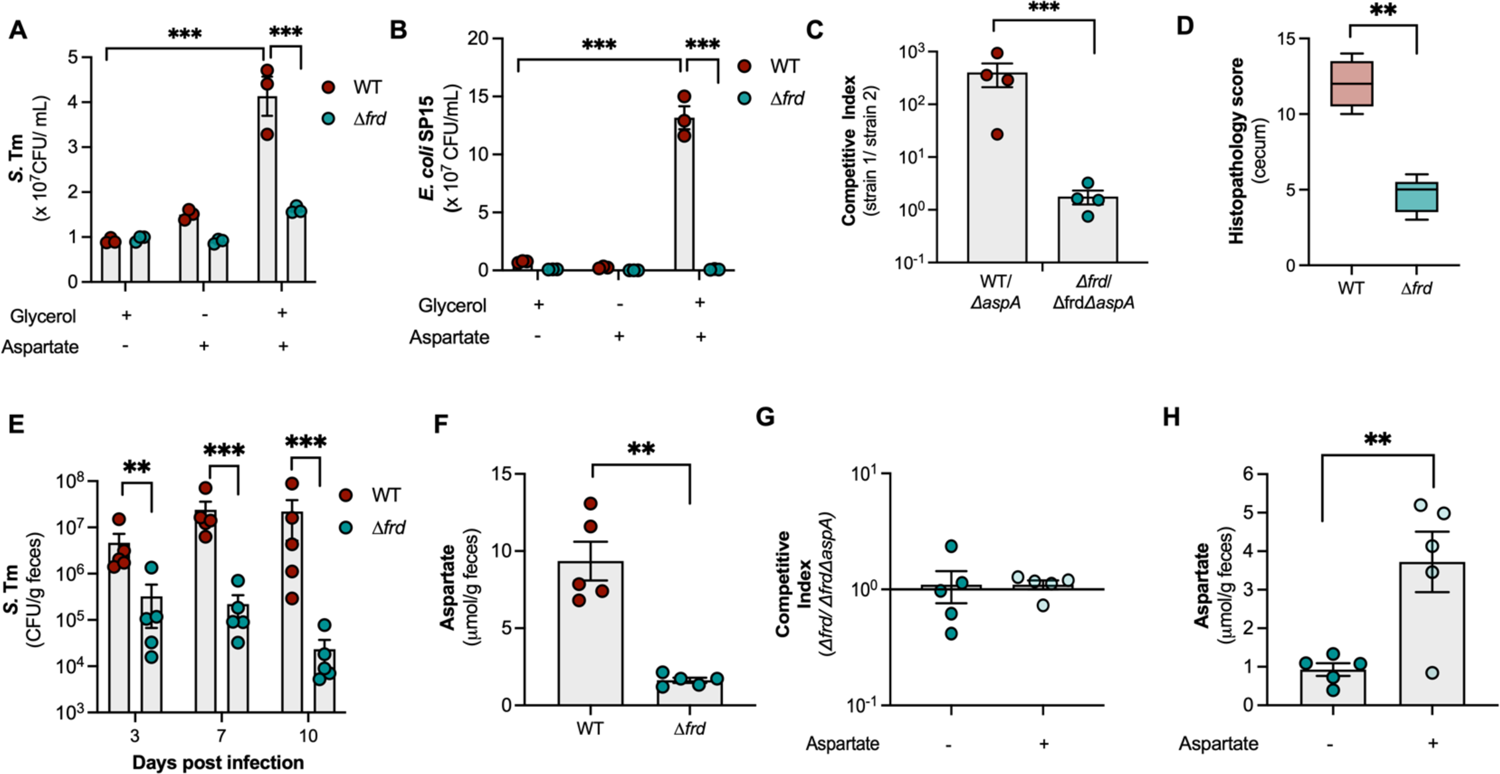
Increased aspartate availability supports fumarate respiration by pathogenic Enterobacteriaceae in the inflamed gut. (A-B) Wild-type *S.* Tm (A) and *Escherichia coli* SP15 (B) and isogenic Δ*frd* mutants were grown in NCE minimal media supplemented or not with glycerol (40 mM) and L-aspartate (30 mM) for 18 hours under anaerobic conditions. (C) Female CBA/J mice were intragastrically infected with 10^9^ CFU of a 1:1 ratio of wild-type *S.* Tm (WT) and isogenic Δ*aspA* mutant or *S.* Tm Δ*frd* and isogenic Δ*frd* Δ*aspA* mutant for ten days. Competitive index in fecal samples was determined at day ten post-infection. (D) Combined histopathology score of pathological lesions in the cecum of mice from (C) at ten days post-infection (n=4). (E) Female CBA/J mice were intragastrically infected with 10^9^ CFU of wild-type *S*. Tm (WT) or isogenic Δ*frd* mutant for ten days. Bacterial numbers in fecal samples were determined at day ten post-infection. (F) Fecal aspartate levels from mice in (E) measured at day ten post-infection. (G) Female CBA/J mice were intragastrically infected with 10^9^ CFU of a 1:1 ratio of *S*. Tm Δ*frd* and Δ*frd* Δ*aspA* isogenic mutant for ten days. A subset of mice received L-aspartate (0.4%) supplementation in their drinking water for the duration of the experiment (Aspartate +). (H) Fecal aspartate levels from mice in (E) measured at day ten post-infection. For *in vitro* experiments, each dot represents one biological replicate (average of triplicate technical replicate per biological replicate). For mouse experiments, each dot represents data from one animal (biological replicate). Bars represent mean ± SEM. The boxes in the whisker plot represent the first to third quartiles, and the mean value of the gross pathology scores is indicated by a line. **, p < 0.01; ***, p < 0.001. See also Figure S2.

Our results (**Fig. 4A-B**) and previous reports (Nguyen et al., 2020; Schubert et al., 2021) demonstrate that pathogenic Enterobacteriaceae can use aspartate to support fumarate respiration *in vitro*. However, whether increased aspartate availability supports pathogenic Enterobacteriaceae expansion in the inflamed gut in a fumarate respiration-dependent manner remained unknown. To address this gap in knowledge, we intragastrically infected CBA/J mice with a 1:1 ratio of wild-type *S.* Tm and an isogenic *ΔaspA* mutant or a 1:1 ratio of *Δfrd* and isogenic *Δfrd ΔaspA* mutants and determined the competitive index between strains at day 10 post-infection. Wild-type *S.* Tm was able to significantly outcompete the *ΔaspA* mutant (**Fig. 4C**) when intestinal inflammation is established (**Fig. 4D**). The aspartate-dependent *S.* Tm growth advantage over an aspartate-deficient mutant was abrogated in mice colonized with *Δfrd* and isogenic *Δfrd ΔaspA* mutants (**Fig. 4C**), suggesting that aspartate fuels *S.* Tm fumarate respiration in the inflamed gut.

Next, we sought to investigate the role of chemotaxis towards and transport of exogenous aspartate in supporting fumarate-dependent *S.* Tm respiration during colitis. We intragastrically infected CBA/J mice with a 1:1 ratio of wild-type *S.* Tm and an isogenic *Δtar* mutant or *Δfrd* and isogenic *Δfrd Δtar* (**Fig. S2B**) or 1:1 ratio of wild-type *S.* Tm and an isogenic *ΔdcuA* mutant *Δfrd* and isogenic *Δfrd ΔdcuA* (**Fig. S2C**) mutants and assessed the competitive index between strains at day 10 post-infection. Wild-type *S.* Tm was able to significantly outcompete the *Δtar* mutant (**Fig. 3D and S2B**) and the *ΔdcuA* (**Fig. 3D and S2C**) mutant during colitis. This growth advantage was abolished in mice colonized with *Δfrd* and isogenic *Δfrd Δtar* mutant (**Fig. S2C**) or in mice colonized with *Δfrd* and isogenic *Δfrd ΔducA* mutant. These results revealed the need for *S.* Tm to perform chemotaxis and gain access to exogenous aspartate to support fumarate respiration during infectious colitis.

To confirm that *S*. Tm *Δfrd* mutant is defective in colonizing the intestinal lumen during colitis, we repeated the experiment by performing single infections in the CBA/J model with wild-type *S*. Tm and isogenic *Δfrd* mutant. We determined intestinal colonization by *S*. Tm strains at day three, seven and ten post-infection. Wild-type *S.* Tm growth in the intestinal lumen significantly increased as disease progressed, reaching higher levels at day seven and ten post-infection (**Fig. 4E**). However, *S.* Tm *Δfrd* was defective in colonizing the intestinal lumen (**Fig. 4E**) and in inducing colitis (**Fig 4D**), suggesting that fumarate respiration is necessary for *S.* Tm to thrive in the gut and induce intestinal inflammation. Importantly, the inability of *S.* Tm *Δfrd* mutant to cause intestinal inflammation was not due to an invasion defect, as the *S.* Tm *Δfrd* strain was able to invade intestinal epithelial cells to similar levels as wild-type *S.* Tm *in vitro* (**Fig. S2D)**. The concentration of aspartate was measured in the feces of mice infected with wild-type or *Δfrd* and a significant difference in fecal aspartate levels was observed (**Fig. 4F**) indicating that the decreased ability of *S.* Tm *Δfrd* to colonize the intestinal lumen in our infectious colitis model could be due to decreased levels of aspartate in *Δfrd*-infected mice. To address this concern, we repeated the competitive infection experiment described in **Fig. 4C** and increased intestinal concentrations of aspartate in *Δfrd*-infected mice by providing aspartate (0.4%) in the drinking water throughout the experiment (**Fig. 4G-H**). Aspartate supplementation did not increase the ability of the fumarate. respiration-deficient *S.* Tm strain to outcompete an isogenic *ΔaspA* mutant at ten days post-infection (**Fig. 4G**), despite the increased aspartate levels in the intestinal lumen of *S.* Tm Δ*frd*-infected mice that received aspartate supplementation (**Fig. 4H**). Taken together, our findings suggest that inflammation-mediated increase in aspartate levels is necessary to support fumarate-dependent intestinal expansion of pathogenic Enterobacteriaceae in the inflamed gut.

### Dietary aspartate is not the main source of this amino acid during intestinal inflammation

Previous work suggests that dietary aspartate fuels aspartate-dependent fumarate respiration during early intestinal Enterobacteriaceae infection, before the onset of inflammation (Nguyen et al., 2020; Schubert et al., 2021). However, the source of aspartate (**Fig. 1A and B**) in the inflamed gut is unknown. To assess whether diet-derived aspartate facilitates *aspA*-dependent *S*. Tm expansion during intestinal inflammation, we determined the ability of wild-type *S*. Tm to outcompete an isogenic *ΔaspA* mutant in the intestines of CBA/J mice fed a control (Asp+) and an aspartate-free (Asp-) diet (**Fig. 5A-C**). To our surprise, wild-type *S*. Tm had a significant growth advantage over the *ΔaspA* mutant in both Asp+ and Asp-diet groups (**Fig. 5A**). This phenotype was accompanied by increased aspartate levels in the intestinal lumen of mice fed either diet (**Fig. 5B**) as well as comparable intestinal inflammation in both groups (**Fig. 5C**). Similar results were obtained in DSS-treated mice infected with wild-type *E. coli* SP15 and an isogenic *ΔaspA* mutant (**Fig. 5D-F**). Our findings demonstrate that dietary aspartate is not the main source of this amino acid during infectious and noninfectious colitis and does not contribute to aspartate-dependent Enterobacteriaceae expansion during intestinal inflammation.

**Figure 5.**
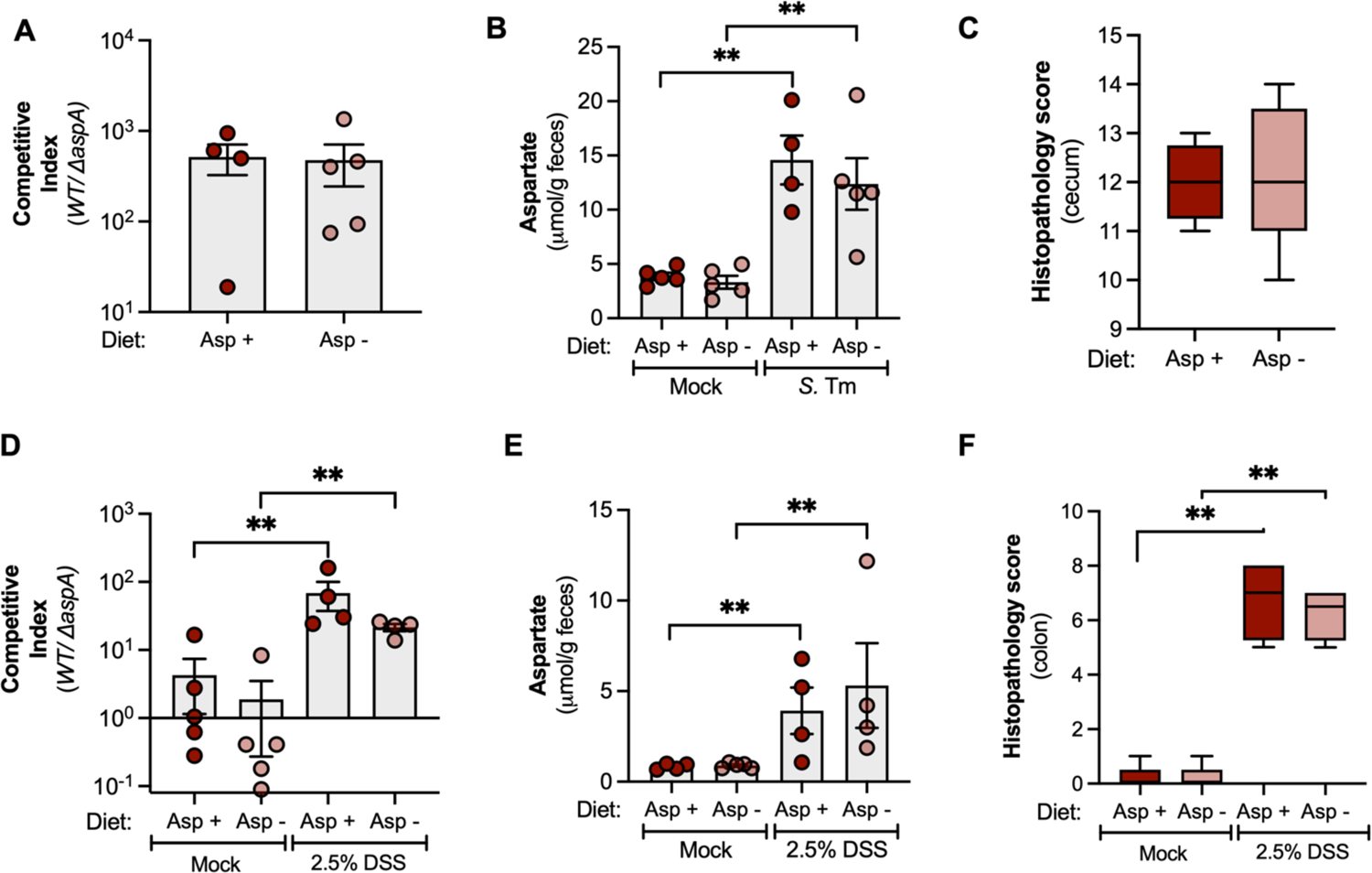
Dietary aspartate is not a major source of aspartate for pathogenic Enterobacteriaceae in the inflamed gut. (A) Female CBA/J mice received a control (Asp +) or an aspartate-free diet (Asp −) for the duration of the experiment. Mice were intragastrically infected with 10^9^ CFU of a 1:1 ratio of wild-type *S*. Tm (WT) and isogenic Δ*aspA* for ten days. Competitive index in fecal samples was determined at day ten post-infection. (B) Fecal aspartate levels from mice in (A) measured at day ten post-infection. (C) Combined histopathology score of pathological lesions in the cecum of mice from (A) at ten days post-infection (n=5). (D) Female C57Bl6/J mice received a control (Asp +) or an aspartate-free diet (Asp −) for the duration of the experiment. Female C57BL/6J mice were intragastrically infected with 10^9^ CFU of a 1:1 ratio of wild-type *E. coli* SP15 (WT) and isogenic Δ*aspA* mutant. One day after infection mice were treated with 2.5% Dextran Sodium Sulfate (DSS) in their drinking water for seven days. Competitive index in fecal samples was determined at day eight post-infection. (E) Fecal aspartate levels from mice in (D) measured at day eight post-infection. (F) Combined histopathology score of pathological lesions in the cecum of mice from (D) at eight days post-infection (n=5). Each dot represents data from one animal (biological replicate). Bars represent mean ± SEM. The boxes in the whisker plot represent the first to third quartiles, and the mean value of the gross pathology scores is indicated by a line. **, p < 0.01.

### Microbiota-derived aspartate fuels pathogenic Enterobacteriaceae expansion in the inflamed gut

Next, we sought to assess whether the host or the gut microbiota were the source of aspartate in the inflamed gut. To test whether the host response is the major source of aspartate during intestinal inflammation, we measured aspartate levels in fecal contents of germ-free mouse models of infectious and noninfectious colitis. We used germ-free Swiss-Webster (SW) mice, a mouse strain that is susceptible to *S.* Tm-induced gastroenteritis (Gillis et al., 2018; Shelton et al., 2021), and DSS-induced colitis (Hughes et al., 2017). Importantly, mock-treated germ-free SW mice had very low aspartate levels in their feces (**Fig. S3A**). To induced infectious colitis in our germ-free SW mouse model, we intragastrically infected mice with wild-type *S.* Tm for three days. We were unable to carry the experiment beyond a three-day time-point as germ-free mice are highly susceptible to *S.* Tm induced intestinal inflammation (Shelton et al., 2021). To induce colitis independently of a pathogen (noninfectious colitis), we treated germ-free SW mice with 2.5% dextran sodium sulfate (DSS) in the drinking water for seven days. Interestingly, we observed no increase in aspartate levels in fecal samples from *S*. Tm-infected gem-free mice (**Fig. S3A**) and DSS-treated germ-free mice (**Fig. S3A**) when compared to mock-treated animals, despite the development of intestinal inflammation (**Fig. S3B**). These results provide the initial evidence that the host inflammatory response alone is not the main contributor to the increased levels of luminal aspartate during colitis.

To determine whether pathogenic Enterobacteriaceae rely on *aspA*-dependent aspartate utilization *in vivo* in the absence of a resident microbiota, we infected germ-free SW mice with 1:1 ratio of wild-type *S.* Tm and *ΔaspA* mutant and assessed the competitive index between strains three days post infection. *S.* Tm infected germ-free mice developed intestinal inflammation comparable to that observed in conventional mice (**Fig. S3B**). Wild-type *S.* Tm was unable to outcompete the isogenic *ΔaspA* mutant in this model (**Fig. 6A**). Similar results were obtained in DSS-treated germ-free SW mice infected with wild-type *E. coli* SP15 and an isogenic Δ*aspA* mutant (**Fig. 6C**). Our findings suggest that host-derived aspartate does not support the growth of pathogenic Enterobacteriaceae during colitis, raising the possibility that the resident microbiota is supporting aspartate-dependent pathogen-expansion during intestinal inflammation.

**Figure 6.**
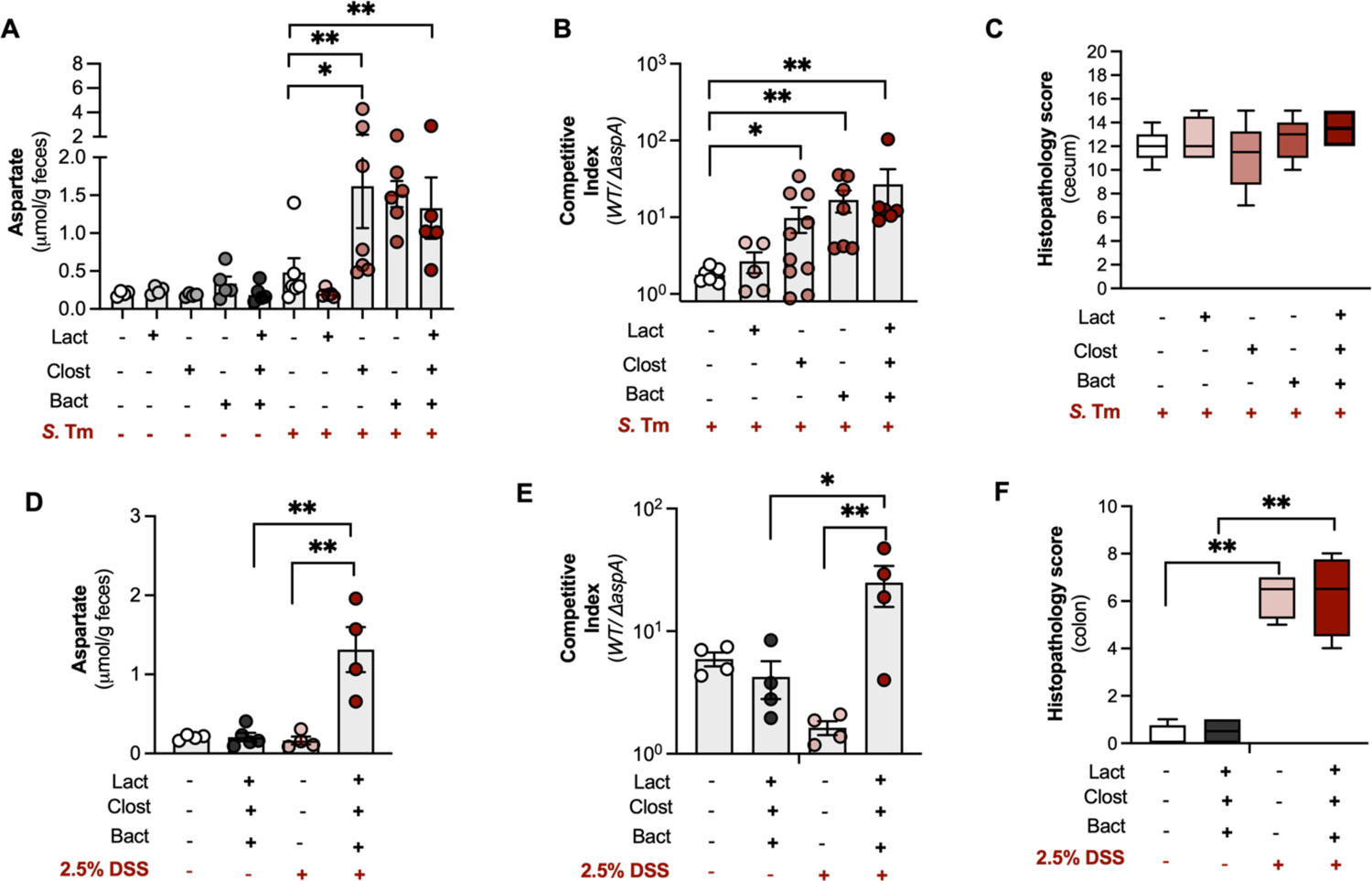
Microbiota-derived aspartate supports pathogenic Enterobacteriaceae expansion in the inflamed gut. (A-C) Female and male Swiss-Wester (SW) germ-free mice were pre-colonized with individual species or a combination of gut commensal microbes for five days. Bact: *Bacteroides caecimuris;* Clost: a combination of *Blautia coccoides, Clostridium cochlearium and Clostidium sporogenes;* Lact: *Lactobacillus reuteri.* After five days, mice were intragastrically infected with 10^9^ CFU of a 1:1 ratio of wild-type *S*. Tm (WT) and isogenic Δ*aspA* for three days. (A) Fecal aspartate levels measured at three days after *S*. Tm infection. (B) Competitive index of *S*. Tm strains in fecal samples was determined at day three post-infection. (C) Combined histopathology score of pathological lesions in the cecum of mice from (B) at three days post-infection (n=5-10). (D-F) Female and male SW germ-free mice were pre-colonized with individual species or a combination of gut commensal microbes for five days. Bact: *Bacteroides caecimuris;* Clost: a combination of *Blautia coccoides, Clostridium cochlearium and Clostidium sporogenes;* Lact: *Lactobacillus reuteri.* After five days, mice were intragastrically infected with 10^9^ CFU of a 1:1 ratio of wild-type *E. coli* SP15 (WT) and isogenic Δ*aspA* mutant. One day after infection mice were treated with 2.5% Dextran Sodium Sulfate (DSS) in their drinking water for seven days. Competitive index in fecal samples was determined at day eight post-infection(D) Fecal aspartate levels measured at seven days after DSS treatment. (E) Competitive index of *E. coli* strains in fecal samples was determined at day eight post-infection. (F) Combined histopathology score of pathological lesions in the cecum of mice from (E) at eight days post-infection (n=4). Each dot represents data from one animal (biological replicate). Bars represent mean ± SEM. The boxes in the whisker plot represent the first to third quartiles, and the mean value of the gross pathology scores is indicated by a line. *, p<0.05; **, p < 0.01. See also Figure S3.

To further explore the hypothesis that the resident microbiota is the major source of aspartate during intestinal inflammation, we pre-colonized germ-free SW mice with commensal microbes representing the most abundant commensal bacteria classes in the mouse gut (**Fig. S3C**) (Winter et al., 2010) and repeated the competitive infection experiments described above. Interestingly, the increased aspartate levels as well as the ability of wild-type *S*. Tm to outcompete the *ΔaspA* mutant were restored in the intestinal lumen of germ-free mice pre-colonized with constituents from the classes Bacteroidia (*Bacteroides caecimuris*) or Clostridia (a combination of *Clostridium clostridioforme*, *Clostridium innocuum*, *Clostridioides mangenotti*, *Clostridium cochlearium* and *Clostridium sporogenes*), but not when mice were pre-colonized with constituents of the class Bacilli (*Lactobacillus reuteri*). Importantly, differences in the competitive advantage between wild-type *S.* Tm and an *ΔaspA* mutant were not due to differences in intestinal inflammation between groups (**Fig. 6C**). Similar results were obtained in DSS-treated germ-free SW mice pre-colonized with members of the microbiota and infected with wild-type *E. coli* SP15 and an isogenic *ΔaspA* mutant (**Fig. 6D-F**). Noninfectious intestinal inflammation induced increased aspartate levels in the intestinal lumen (**Fig. 6D**) and a competitive advantage *of E. coli* SP15 over an isogenic *ΔaspA* mutant (**Fig. 6E**) only in mice pre-colonized with a defined microbiota, despite similar levels of intestinal inflammation between groups (**Fig. 6F**). Taken together, our findings demonstrate the role for microbiota-derived aspartate in driving respiration-dependent expansion of pathogenic Enterobacteriaceae in the inflamed gut.

## DISCUSSION

The gut microbiota occupies major nutrient niches to limit pathogen expansion in the gastrointestinal tract (colonization resistance) (Shealy et al., 2021). In particular, commensal microbes provide colonization resistance by sequestering amino acids (Caballero-Flores et al., 2020), reducing the pool of a key nutrient source for pathogenic Enterobacteriaceae expansion in the intestinal lumen (Kitamoto et al., 2020; Nguyen et al., 2020; Schubert et al., 2021). Thus, enteric pathogens must develop strategies to alter the gut environment in a way that allows for them to gain access to amino acids that are necessary for successful intestinal colonization (Nguyen et al., 2020; Schubert et al., 2021). In this study we report that intestinal inflammation causes increased levels of the amino acid aspartate in the intestinal lumen, in a microbiota-dependent manner. Our data show that *S.* Tm and *pks*+ *E. coli* (*E. coli* SP15) can use this new source of exogenous aspartate to support fumarate-dependent anaerobic respiration in the inflamed gut. This leads to an advantage *in vivo* as *S.* Tm and *E. coli* SP15 strains that can transport aspartate and perform aspartate-dependent fumarate respiration outcompete isogenic strains that cannot utilize this amino acid. Collectively, we describe a previously unexplored mechanism by which pathogenic Enterobacteriaceae exploit the gut microbiota to acquire the amino acid aspartate and support aspartate-dependent anaerobic respiration in the inflamed gut (Fig. S4).

Recent reports identify dietary aspartate as a potential source of fumarate-dependent respiration during early intestinal Enterobacteriaceae infection, before the onset of inflammation (Nguyen et al., 2020; Schubert et al., 2021). However, in this study we show that dietary aspartate does not contribute to *S.* Tm and *E. coli* SP15 expansion in the inflamed gut. We identified the gut microbiota, especially members of the class Bacteroidia and Clostridia, as the major source of aspartate during intestinal inflammation. It remains unclear how inflammation can induce the release of amino acids by the gut microbiota. One possibility is that the innate immune response elicited by *S*. Tm infection or noninfectious colitis stimulates the production of toxic reactive oxygen species (ROS) and antimicrobial peptides, which can induce bacterial lysis of microbiota constituents (Li et al., 2020; Lin and Zhang, 2017; Takiishi et al., 2017). Alternatively, the introduction of oxygen as a function of inflammation (Rivera-Chávez et al., 2016) may alter key metabolic pathways leading to inadvertent secretion of aspartate by the microbiota in response to an adaptation to the loss of intestinal anaerobiosis (Lin and Zhang, 2017; Takiishi et al., 2017). Understanding the mechanism of amino acid release by the gut microbiota as a consequence of the host inflammatory response may have significant implications in the pathogenesis of infectious and noninfectious colitis and will be an important area of future research.

Commensal Enterobacteriaceae play a crucial role in colonization resistance against *S*. Tm and other pathogenic members of this family (Brugiroux et al., 2016; Litvak et al., 2019; Velazquez et al., 2019; Wotzka et al., 2019). In particular, commensal Enterobacteriaceae compete for carbon sources (Eberl et al., 2021) and electron acceptors generated by the host response (Litvak et al., 2019; Velazquez et al., 2019) during early infection. However, the metabolic interactions between commensal and pathogenic Enterobacteriaceae during the late stages of colitis (once intestinal inflammation is established) remain unclear. The gene encoding aspartate ammonia-lyase (*aspA*) is conserved amongst commensal *E. coli* species and non-pathogenic members of the Enterobacteriaceae family can perform aspartate-dependent fumarate respiration (Schubert et al., 2021) (data not shown). Such findings suggest that not only pathogenic Enterobacteriaceae but also commensal species belonging to this family may have access to this novel source of electron acceptors to expand in the inflamed gut. Thus, the mechanisms used by *S.* Tm to outcompete commensal Enterobacteriaceae aspartate utilization and gain access to this key resource remain to be elucidated.

In conclusion, our findings provide a fundamental advance in understanding how pathogenic Enterobacteriaceae have evolved to utilize microbiota-derived metabolites to thrive in the inflamed gut. The emerging picture is that enteric pathogens rely on microbiota-derived amino acids as a source of electron acceptors for growth via anaerobic respiration during intestinal inflammation. Our findings may aid in the development of novel treatment strategies focused on pathogen metabolic adaptations for infectious and noninfectious gastroenteritis.

## Acknowledgments

W.Y. was supported by the Basic Science Research Program through the National Research Foundation of Korea (N.R.F.) by the Ministry of Education 2020R1A6A3A03037326. N.G.S. was supported by NIH T32 Training Grant T32ES007028-46 and GT15104 from the Howard Hughes Medical Institute through the James H. Gilliam Fellowships for Advanced Study program. C.D.S. was supported by Dorothy Beryl and Theodore Roe Austin Pathology Research Fund, NIH T32 Training Grant T32AI112541 and NIH Ruth L. Kirschstein Predoctoral Individual Fellowship 1F31AI161882-01A1. N.J.F. was supported by NIH T32 Training Grant T32DK007673. Work in M.X.B.’s laboratory was supported by V Scholar grant V2020-013 from The V Foundation for Cancer Research, Vanderbilt Digestive Disease Pilot and Feasibility grant P30 058404, A.C.S. Institutional Research Grant IRG-19-139-59, VICC GI SPORE grant P50CA236733, United States-Israel Binational Science Foundation grant 2019136 and Vanderbilt Institute for Clinical and Translational Research Grant VR53102.

## Author contributions

W.Y., J.K.Z., and M.X.B. designed and conceived the study. W.Y., J.K.Z., N.G.S., T.P.T., J.D.T., C.D.S., N.J.F., and E.E.O. performed all experiments. All authors contributed to the data analysis and preparation of the manuscript. M.X.B. secured funding for the study.

## MATERIAL AND METHODS

### RESOURCES AVAILABILITY

#### Lead contact

Further information and requests for resources and reagents should be directed to and will be fulfilled by the lead contact, Mariana X. Byndloss.

#### Materials availability

All unique reagents generated in this study are available from the lead contact without restriction.

#### Data and code availability

Data reported in this paper will be shared by the lead contact upon request. This paper does not report original code. Any additional information required to reanalyze the data reported in this paper is available from the lead contact upon request.

### EXPERIMENTAL MODEL AND SUBJECT DETAILS

#### Mouse husbandry

All animal experiments in this study were approved by the Institutional Animal Care and Use Committee at the Vanderbilt University Medical Center. Female CBA and C57BL/6 mice were obtained from the Jackson Laboratory. Germ-Free Swiss Webster (SW) mice were bred in house and maintained in specific pathogen-free facilities at Vanderbilt University Medical Center. A subset of germ-free mice was colonized with a defined microbiota to obtain the SW mice used in this study. The age at the beginning of the experiment was 6-7 weeks old for all CBA, C57BL/6 and SW mice. Conventional mice were housed in individually ventilated cages with ad *libitum* access to water and chow (Envigo Global 16% Protein Rodent Diet). Germ-free SW mice were housed in sterile positive pressure cages (Allentown) with ad *libitum* access to sterile water and chow.

For the germ-free SW mice, unless indicated otherwise in the figure legend, both male and female mice were analyzed and no significant sex-specific differences were noted. Both sexes were equally represented in each experimental group. Animals were randomly assigned into cages and treatment groups 3 days prior to experimentation. Unless stated otherwise, a minimum of 5 mice were used based on variability observed in previous experiments. All mice were monitored twice per day and cage bedding changed every two weeks. At the end of the experiments, mice were humanely euthanized using carbon dioxide inhalation. Animals that had to be euthanized for humane reasons prior to reaching the predetermined time point were excluded from the analysis.

#### Bacterial strains

*S*. Typhimurium IR715 and *E. coli* SP15 strains were routinely grown aerobically at 37°C in LB broth (10 g/L tryptone, 5 g/L yeast extract, 10 g/L sodium chloride) or on LB agar plates (10 g/L tryptone, 5 g/L yeast extract, 10 g/L sodium chloride, 15 g/L agar). For growth under anaerobic conditions, No-carbon-E (NCE) minimal medium (28 mM KH_2_PO_4_, 28 mM K_2_HPO_4_×3H_2_O and 16 mM NaNH_4_HPO_4_×4H_2_O) supplemented with 1 mM MgSO_4_, 0.1% Casamino acids and 1% Vitamin and Trace mineral solutions (American Type Culture Collection) was used. As a sole carbon source, glycerol (40 mM) was added to the bacterial culture, and L-aspartate (30 mM) was also used as a source of fumarate for anaerobic fumarate respiration. All *S*. Typhimurium IR715 and *E. coli* SP15 strains were incubated at 37°C in an anaerobic chamber (0% oxygen). When appropriate, agar plates and media were supplemented with 30 μg/mL chloramphenicol (Cm), 100 μg/mL carbenicillin (Carb), 50 μg/mL kanamycin (Km), or 50 μg/mL nalidixic acid (Nal). Defined-microbiota (DM) strains were cultured in an anaerobic chamber (85% nitrogen, 10% hydrogen, 5% carbon dioxide, Coy Lab Products). *Bacteroides caecimuris* strain (Key Resources Table) was routinely cultured on blood agar plates (37 g/L brain heart infusion medium, 15 g/L agar, 50 mM sheep blood). *Lactobacillus reuteri* strain (Key Resources Table) was routinely cultured on MRS agar plates (55 g/L MRS medium, 15 g/L agar). *Clostridium clostridioforme*, *Clostridium clostridioforme*, *Clostridioides mangenotti*, *Clostridium cochlearium* and *Clostridium sporogenes* strains (Key Resources Table) were routinely cultured on BHI agar plates (37 g/L brain heart infusion medium, 15 g/L agar).

## METHODS DETAILS

### Construction of plasmids and bacterial strains

#### Construction of Salmonella enterica serovar Typhimurium mutants

For the *S*. Typhimurium IR715 Δ*tar*, Δ*aspA*, Δ*dcuA* or Δ*frdABCD* deletion mutants, the upstream and downstream regions of approximately 0.5 kb flanking the *tar*, *aspA*, *dcuA* or *frdABCD* genes were amplified by PCR using primers IR715-tar-del-F1/IR715-tar-del-R1 and IR715-tar-del-F2/ IR715-tar-del-R2, IR715-aspA-del-F1/IR715-aspA-del-R1 and IR715-aspA-del-F2/IR715-aspA-del-R2, IR715-dcuA-del-F1/IR715-dcuA-del-R1 and IR715-dcuA-del-F2/IR715-dcuA-del-R2 or IR715-frdABCD-del-F1/IR715-frdABCD-del-R1 and IR715-frdABCD-del-F2/IR715-frdABCD-del-R2 from the *S*. Typhimurium IR715 wild-type genome. The pRDH10 suicide plasmid was digested with SalI and assembled with the fragments of each gene using the Gibson Assembly Master Mix (NEB) to form plasmid pWJ39 (pRDH10::Δ*tar*), pWJ40 (pRDH10::Δ*aspA*), pWJ41 (pRDH10::Δ*dcuA*) or pWJ51 (pRDH10::Δ*frdABCD*).

Recombinant plasmids pWJ39, pWJ40, pWJ41 or pWJ51 were transformed into *E. coli* S17-1 λ*pir* and conjugated into *S*. Typhimurium IR715 wild-type. Clones that had integrated the suicide plasmid were subjected to sucrose counter-selection and a colony that was sucrose resistant and Cm^S^ was verified by PCR to be *S*. Typhimurium IR715 Δ*tar*, Δ*aspA*, Δ*dcuA* or Δ*frdABCD* deletion mutants.

Recombinant plasmids pWJ39, pWJ40 or pWJ41 were transformed into *E. coli* S17-1 λ*pir* and conjugated into *S*. Typhimurium IR715 Δ*frdABCD* mutant (WJ28) using *E. coli* S17-1 λ*pir* as the donor strain. Clones that had integrated the suicide plasmid were subjected to sucrose counter-selection and a colony that was sucrose resistant and Cm^S^ was verified by PCR to be *S*. Typhimurium IR715 Δ*frdABCD* Δ*tar*, Δ*frdABCD* Δ*aspA* or Δ*frdABC*D Δ*dcuA* mutants.

Recombinant plasmids pWJ39, pWJ40, pWJ41 or pWJ51 were transformed into *E. coli* S17-1 λ*pir* and conjugated into *S*. Typhimurium IR715 Δ*invA* Δ*spiB* mutant (SPN487) using *E. coli* S17-1 λ*pir* as the donor strain. Clones that had integrated the suicide plasmid were subjected to sucrose counter-selection and a colony that was sucrose resistant and Cm^S^ was verified by PCR to be *S*. Typhimurium IR715 Δ*invA* Δ*spiB* Δ*tar*, Δ*invA* Δ*spiB* Δ*aspA*, Δ*invA* Δ*spiB* Δ*dcuA* or Δ*invA* ΔspiB Δ*frdABCD* mutants. The Δ*phoN*::Tn*10d*-Cm^R^ and Δ*phoN*::Km^R^ mutation were transduced by phage P22 HT *int-105* (Schmieger, 1972) from FF176 and AJB715 respectively. Primers and plasmids used to construct the bacterial strains are listed in Table S1 and S2 respectively.

#### Construction of Escherichia coli SP15 mutants

To delete *aspA* or *frdABCD* from the *E. coli* SP15 parent strain, lambda red recombination was used. The Cm^R^ cassette was amplified from pKD3 using the SP15-aspA-del-F1/SP15-aspA-del-R1 primers and the Kan^R^ cassette was amplified from pKD13 using the SP15-frdABCD-del-F1/SP15-frdABCD-del-R1 primers. The resulting PCR product was integrated into the *aspA* or *frdABCD* region in a wild-type *E. coli* SP15 strain containing the plasmid pKD46, followed by the selection of Δ*aspA*::Cm^R^ or Δ*frdABCD*::Kan^R^ mutants. The Cm^R^ or Kan^R^ cassette was removed using the plasmid pCP20 (Datsenko and Wanner, 2000). Primers and plasmids used to construct the bacterial strains are listed in Table S1 and S2 respectively.

#### *In vitro* bacterial growth assays

Growth assays were performed in No-carbon-E (NCE) minimal medium (28 mM KH_2_PO_4_, 28 mM K_2_HPO_4_×3H_2_O and 16 mM NaNH_4_HPO_4_×4H_2_O) supplemented with 1 mM MgSO_4_, 0.1% Casamino acids and 1% Vitamin and Trace mineral solutions (American Type Culture Collection) in the presence or absence of glycerol (40 mM) as a sole carbon source and/or L-aspartate (30 mM) as a source of fumarate. For anaerobic growth experiments, media was placed in the anaerobic chamber 48 h prior to inoculation. Overnight aerobic cultures of *S*. Tm or *E. coli* strains were harvested, washed in PBS, and resuspended in NCE media. Wild-type *S*. Tm or *E. coli* was then added to anaerobic media containing carbon source (40 mM glycerol) and/or a source of fumarate (30 mM L-aspartate), which is used as an electron acceptor, at a final concentration of 1 × 10^4^ CFU/mL. Bacterial growth was determined after 18 h by spreading serial ten-fold dilutions onto LB agar plates containing the appropriate antibiotics.

To measure anaerobic growth of wild-type and mutant strains of *S*. Tm and *E. coli* with glycerol and/or L-aspartate, overnight cultures of each strain were harvested, washed in PBS, and resuspended in NCE media. In a 96 well plate, strains were added to NCE media containing 40 mM glycerol or 30 mM L-aspartate, or a combination of both at a final OD_600_=0.1. OD_600_ was measured for 24 h using the Epoch 2 plate reader (Bio-Tek). *In vitro* bacterial growth assays were performed in triplicate with different colonies.

### Bacterial RNA isolation and quantitative real-time PCR

To determine bacterial gene expression *in vitro*, NCE minimal medium (28 mM KH_2_PO_4_, 28 mM K_2_HPO_4_×3H_2_O and 16 mM NaNH_4_HPO_4_×4H_2_O) supplemented with 1 mM MgSO_4_, 0.1% Casamino acids and 1% Vitamin and Trace mineral solutions (American Type Culture Collection) was inoculated with *S*. Typhimurium IR715 strains and cultured at 37°C under anaerobic condition. Glycerol (40 mM) was provided as a sole carbon/energy source and L-aspartate (30 mM) was added as a source of fumarate for anaerobic fumarate respiration. Total RNA was extracted from the bacterial cells grown to the early-log phase (4 h p.i.) using SurePrep^TM^ TrueTotal^TM^ RNA Purification Kit (Fisher BioReagents) or Total RNA Purification Plus Micro Kit (Norgen). cDNA was generated by iScript gDNA Clear cDNA Synthesis Kit (Bio-Rad). Real-time PCR was performed using iQ SYBR Green Supermix (Bio-Rad). Data was acquired in a CFX384 Real-Time System (Bio-Rad) and analyzed using the comparative Ct method. Target gene transcription of each sample was normalized to 16S rRNA mRNA levels. Primers used to determine bacterial gene expression in this experiment are listed in Table S3.

### Microbiota analysis

Composition of the microbiota was analyzed as described previously (Barman et al., 2008). Groups of CBA mice were mock-infected or infected with the *S*. Typhimurium wild-type (IR715), and samples were collected ten days after infection. The bacterial genomic DNA was extracted using the PowerSoil DNA Isolation Kit (Mo-Bio) according to the manufacturer’s instructions. Bacterial DNA was diluted 1:10 and a 0.04 mL served as the template for SYBR-green based real-time PCR using the primers listed in Table S3. Gene copy number for each sample was determined using standard curves generated from fragments of bacterial 16S rRNA genes (*Eubacteria*: *R. productus* [ATCC 27340D], *Clostridiales: R. productus* [ATCC 27340D], *Lactobacillales/Bacillales*: *L. acidophilus* [ATCC 4357D], *Bacteroidetes/Actinobacteria*: *B. fragilis* [ATCC 25285D], *Enterobacteriaceae*: *E. coli* [ATCC TOP10]) cloned into pCR2.1 (TOPO TA cloning kit, Invitrogen) as templates. The fraction of each population was calculated by dividing the gene copy number of each population by the total gene copy number determined using the universal primers.

### Growth of defined-microbiota (DM) strains

Porcine mucin was dissolved in 1x NCE salts at a final concentration of 0.5% (w/v). Mucin broth was inoculated with a fresh colony of *Bacteroides caecimuris* and incubated under anaerobic conditions for 72 h at 37°C. MRS broth was inoculated with a fresh colony of *Lactobacillus reuteri* and incubated under anaerobic conditions for 72 h at 37°C. BHI broth was inoculated with a fresh colony of *Clostridium clostridioforme*, *Clostridium clostridioforme*, *Clostridioides mangenotti*, *Clostridium cochlearium* or *Clostridium sporogenes* strain and incubated under anaerobic conditions for 72 h at 37°C.

### Animal experiments

#### Animal models of *S*. Typhimurium-induced colitis

*CBA mouse model:* CBA mice were infected with either 0.1 mL of LB broth (mock-infected) or *S*. Typhimurium IR715 strains in LB broth. Groups (n=4-5) of 6-7 weeks old CBA mice were intragastrically infected with 1 x 10^9^ CFU for single strain and competitive infection experiments. Fecal samples were collected at days 1, 3 and 7 after infection for bacterial plating. Ten days after infection, samples for histopathology, colonic luminal content and cecal content for *S.* Tm enumeration and aspartate measurements were collected. *S.* Tm numbers were determined by plating serial ten-fold dilutions onto LB agar containing the appropriate antibiotics. For the aspartate-free diet experiments, Groups (n=5) of 6-7 weeks old CBA mice were fed control diet A20073101 (Research Diets) or aspartate-free diet A20073102 (Research Diets) beginning 48h before *S.* Tm infection and were kept on the diets throughout the experiment. Remaining experiment procedures were performed as described above.

*Germ-Free Swiss Webster (SW) mice:* Groups (n=5) of 6-7 weeks old SW germ-free mice were intragastrically inoculated with 3 x 10^9^ CFU of *Bacteroides caecimuris*, *Lactobacillus reuteri*, or a combination of *Clostridium clostridioforme*, *Clostridium innocuum*, *Clostridioides mangenotti*, *Clostridium cochlearium* and/or *Clostridium sporogenes*. A subset of mice was not pre-colonized and remained germ-free. After 7 days, pre-colonized and germ-free control animals were infected with 1 x 10^9^ *Salmonella enterica* serovar Typhimurium IR715 strains. Three days after infection, samples for histopathology, colonic luminal content and cecal content for *S.* Tm enumeration and aspartate measurements were collected. *S.* Tm numbers were determined by plating serial ten-fold dilutions onto LB agar containing the appropriate antibiotics.

#### Animal models of DSS-induced colitis

*Conventional Mice.* Male and female 6-7 weeks old wild-type C57BL/6 mice were infected with 1 x 10^9^ CFU of a 1:1 mixture of *E. coli* SP15 strains. One day post infection, mice were given 2.5% dextran sulfate sodium (DSS; Alfa Aesar) in their drinking water continuously for 7 days. At 8 days post-infection, samples for histopathology, colonic luminal content and cecal content for *E. coli* SP15 enumeration and aspartate measurements were collected. *E. coli* SP15 numbers were determined by plating serial ten-fold dilutions onto LB agar containing the appropriate antibiotics. For the aspartate-free diet experiments, groups (n=5) of 6-7 weeks old C57Bl/6 mice were fed control diet A20073101 (Research Diets) or aspartate-free diet A20073102 (Research Diets) beginning 48h before *E. coli* SP15 infection and were kept on the diets throughout the experiment. Remaining experiment procedures were performed as described above.

Germ-Free Swiss Webster (SW) mice: Groups (n=5) of 6-7 weeks old germ-free SW mice were intragastrically inoculated with 3 x 10^9^ CFU of Bacteroides caecimuris, Lactobacillus reuteri, or a combination of Clostridium clostridioforme, Clostridium innocuum, Clostridioides mangenotti, Clostridium cochlearium and/or Clostridium sporogenes. A subset of mice were not pre-colonized and remained germ-free. After 7 days, pre-colonized and germ-free control animals were infected with with 1 x 10^9^ CFU of a 1:1 mixture of E. coli SP15 strains. One day post infection, mice were given 2.5% dextran sulfate sodium (DSS; Alfa Aesar) in their drinking water continuously for 7 days. Samples were collected at 8 days post-infection. At 8 days post-infection, samples for histopathology, colonic luminal content and cecal content for E. coli SP15 enumeration and aspartate measurements were collected. E. coli SP15 numbers were determined by plating serial ten-fold dilutions onto LB agar containing the appropriate antibiotics.

### Caco-2 cell culture

The human epithelial colorectal adenocarcinoma cell line Caco-2 was obtained from the American Type Culture Collection. Caco-2 cells were routinely maintained in 1x Minimal Essential Media (Gibco), with 10% Fetal Bovine Serum (Gibco), 1x GlutaMax (Gibco), 1x MEM Nonessential Amino Acids (Corning), 1 mM Sodium Pyruvate (Gibco) at 37°C and 5% CO_2_.

### Invasion assay

Caco-2 cells were seeded 2.5 x 10^5^ cells/well and infected with indicated strains at a multiplicity of infection (MOI) of 10. The bacteria and cells were exposed to hypoxic conditions in a humidified hypoxia chamber (0.8% O_2_) while being incubated for 1 h at 37°C in 1x Minimal Essential Media (Gibco), with 10% Fetal Bovine Serum (Gibco), 1x GlutaMax (Gibco), 1x MEM Nonessential Amino Acids (Corning), 1 mM Sodium Pyruvate (Gibco). Each well was washed three times with sterile PBS (KCl at 2.7 mM, KH_2_PO_4_ at 1.8 mM, NaCl at 140 mM, Na_2_HPO_4_ at 10 mM, pH 7.4) to remove extracellular bacteria, and medium containing gentamicin at a concentration of 0.1 mg/mL was added for a 90 min incubation in conditions stated above. After three washes with PBS, the cells were lysed with 1 mL of 1% Triton X-100 and the lysates were transferred to sterile tubes. Tenfold serial dilutions were plated to enumerate intracellular bacteria.

### Aspartate measurement

The aspartate concentration in cecal contents of mice used in this study was assessed using an Aspartate Colorimetric/Fluorometric Assay Kit (BioVision) according to the manufacturer’s instructions.

### Histopathology scoring

Cecal and colonic tissue was fixed in 10% phosphate-buffered formalin for 48 h and embedded in paraffin. Sections (5 µm) were stained with hematoxylin and eosin. Stained sections were blinded and evaluated by a veterinary pathologist according to the criteria listed in Table S4.

### Quantification and statistical analysis

Statistical data analysis was performed using Graphpad PRISM. Fold changes of ratios (bacterial competitive index and mRNA levels), and bacterial numbers were transformed logarithmically prior to statistical analysis. An unpaired Student’s t test was used on the transformed data to determine whether differences between groups were statistically significant (p < 0.05). When more than two treatments were used, statistically significant differences between groups were determined by one-way ANOVA followed by Tukey’s HSD test (between > 2 groups). Significance of differences in histopathology was determined by a one-tailed non-parametric test (Mann-Whitney).

## KEY RESOURCES TABLE

| REAGENT or RESOURCE | SOURCE | IDENTIFIER |
| --- | --- | --- |
| **Bacterial and virus strains** |  |  |
| S. Typhimurium, IR715 ATCC 14028 NaI<sup>R</sup> | Stojiljkovic et al., 1995 | IR715 |
| S. Typhimurium, IR715 $\Delta phoN::Tn10d$-Cm<sup>R</sup> | Faber et al., 2016 | FF176 |
| S. Typhimurium, IR715 $\Delta phoN::Km^R$ | Kingsley et al., 2003 | AJB715 |
| S. Typhimurium, IR715 $\Delta tar \Delta phoN::Km^R$ | This study | WJ65 |
| S. Typhimurium, IR715 $\Delta aspA \Delta phoN::Km^R$ | This study | WJ66 |
| *S. Typhimurium*, IR715 Δ*dcuA*<br>Δ*phoN*::Km<sup>R</sup> | This study | WJ57 |
| *S. Typhimurium*, IR715 Δ*frdABCD*<br>Δ*phoN*::Km<sup>R</sup> | This study | WJ29 |
| *S. Typhimurium*, IR715 Δ*frdABCD* | This study | WJ28 |
| *S. Typhimurium*, IR715 Δ*frdABCD*<br>Δ*phoN*::Tn10d-Cm<sup>R</sup> | This study | WJ30 |
| *S. Typhimurium*, IR715 Δ*frdABCD* Δ*tar*<br>Δ*phoN*::Km<sup>R</sup> | This study | WJ67 |
| *S. Typhimurium*, IR715 Δ*frdABCD* Δ*aspA*<br>Δ*phoN*::Km<sup>R</sup> | This study | WJ68 |
| *S. Typhimurium*, IR715 Δ*frdABCD* Δ*dcuA*<br>Δ*phoN*::Km<sup>R</sup> | This study | WJ58 |
| *S. Typhimurium*, IR715 Δ*invA* Δ*spiB* | Rivera-Chávez et al.,<br>2013 | SPN487 |
| *S. Typhimurium*, IR715 Δ*invA* Δ*spiB*<br>Δ*phoN*::Tn10d-Cm<sup>R</sup> | This study | WJ52 |
| *S. Typhimurium*, IR715 Δ*invA* Δ*spiB*<br>Δ*phoN*::Km<sup>R</sup> | This study | WJ111 |
| *S. Typhimurium*, IR715 Δ*invA* Δ*spiB* Δ*tar*<br>Δ*phoN*::Km<sup>R</sup> | This study | WJ125 |
| *S. Typhimurium*, IR715 Δ*invA* Δ*spiB* Δ*aspA*<br>Δ*phoN*::Km<sup>R</sup> | This study | WJ133 |
| *S. Typhimurium*, IR715 Δ*invA* Δ*spiB* Δ*dcuA*<br>Δ*phoN*::Km<sup>R</sup> | This study | WJ135 |
| *S. Typhimurium*, IR715 Δ*invA* Δ*spiB*<br>Δ*frdABCD* Δ*phoN*::Km<sup>R</sup> | This study | WJ143 |
| *E. coli*, S17-1 λ*pir* *zxx*::RP4 2-(Tet<sup>R</sup>::Mu)<br>(Km<sup>R</sup>::Tn7) λ*pir* *recA1* *thi* *pro* *hsdR* (*r*m<sup>+</sup>) | Simon et al., 1983 | S17-1 λ*pir* |
| *E. coli*, DH5α λ*pir*, *F*<sup>-</sup> *endA1* *hsdR17* (*r*m<sup>+</sup>)<br>*supE44* *thi-1* *recA1* *gyrA* *relA1* $\Delta$(*lacZYA*-<br>*argF*)U189 ϕ80*lacZ*ΔM15 λ*pir* | Grant et al., 1990 | DH5α λ*pir* |
| *E. coli*, SP15 | Johnson et al., 2002 | *E. coli* SP15 |
| *E. coli*, SP15 Δ*aspA*::Km<sup>R</sup> | This study | JZ1 |
| *E. coli*, SP15 Δ*frdABCD*::Km<sup>R</sup> | This study | JZ2 |
| *Bacteroides caecimuris* | DSMZ | DSMZ 26085 |
| *Lactobacillus reuteri* | DSMZ | DSMZ 32035 |
| *Clostridium clostridioforme* | DSMZ | DSMZ 26114 |
| *Clostridium innocuum* | DSMZ | DSMZ 26113 |
| *Clostridioides mangenotti* | DSMZ | DSMZ 29133 |
| *Clostridium cochlearium* | DSMZ | DSMZ 29358 |
| *Clostridium sporogenes* | DSMZ | DSMZ 29420 |
| **Chemicals, peptides, and recombinant proteins** |  |  |
| Gibson Assembly Master Mix | NEB | Cat# E2611L |
| LB Broth, Miller (Luria Bertani) | Becton Dickinson | Cat# 244520 |
| LB Agar, Miller (Luria Bertani) | Becton Dickinson | Cat# 244620 |
| MRS Broth (Difco™ Lactobacilli) | Becton Dickinson | Cat# 288130 |
| Bacto Brain Heart Infusion | Becton Dickinson | Cat# 237500 |
| Sheep Blood | Fisher BioReagents | Cat# R54008 |
| Bacto Tryptone | Becton Dickinson | Cat# 211705 |
| L-Aspartic Acid | Fisher Scientific | Cat# BP374-100 |
| Glycerol | Fisher BioReagents | Cat# G33-4 |
| Mucin from porcine stomach, Type II | Sigma | Cat# M2378 |
| Dextran Sulfate Sodium Salt (DSS) | Alfa Aesar | Cat# J63606 |
| Streptomycin Sulfate | Fisher Scientific | Cat# BP910-50 |
| TRI Reagent | Molecular Research | Cat# TR118 |
| iQ SYBR Green Supermix | Bio-Rad | Cat# 1708882 |
| Platinum PCR Supermix High Fidelity | Invitrogen | Cat# 12532-016 |
| PCR SuperMix | Invitrogen | Cat# 10572-014 |
| Acid Casein Peptone | Fisher Scientific | Cat# BP1424-500 |
| Vitamin Supplement | ATCC | Cat# MD-VS |
| Trace Mineral Supplement | ATCC | Cat# MD-TMS |
| Minimum Essential Media (MEM) | Gibco | Cat# 11090081 |
| Fetal Bovine Serum (FBS) | Gibco | Cat# 10500064 |
| GlutaMAX™-I | Gibco | Cat# 35050 |
| MEM Nonessential Amino Acids Solution (NEAA) | Corning | Cat# 25-025 |
| Sodium Pyruvate | Gibco | Cat# 11360070 |
| DPBS | Gibco | Cat# 14190144 |
| HEPES | Gibco | Cat# 15630080 |
| Gentamicin Solution | Sigma | Cat# G1397 |
| **Critical commercial assays** |  |  |
| Aspartate Colorimetric/Fluorometric Assay Kit | BioVision | Cat# K552 |
| PowerSoil DNA Isolation Kit | Mo-Bio | Cat# 12888 |
| SurePrep™ TrueTotal™ RNA Purification Kit | Fisher BioReagents | Cat# BP2800 |
| Total RNA Purification Plus Micro Kit | Norgen | Cat# 48500 |
| iScript gDNA Clear cDNA Synthesis Kit | Bio-Rad | Cat# 1725035 |
| Zymoclean™ Gel DNA Recovery Kit | Zymo Research | Cat# D4002 |
| PureLink™ Quick Plasmid Miniprep Kit | Invitrogen | Cat# K210011 |
| DNeasy PowerSoil Kit | Qiagen | Cat# 12888 |
| **Experimental models: Organisms/strains** |  |  |
| Mouse: C57BL/6J | Jackson Laboratory | Cat# 000664 |
| Mouse: CBA/J | Jackson Laboratory | Cat# 000656 |
| Mouse: Germ-free Swiss Webster | Byndloss Lab | N/A |
| Caco-2 | ATCC | Cat# HTB-37 |
| **Oligonucleotides** |  |  |
| Primers used in this study, see Table S1 | This study | N/A |
| **Recombinant DNA** |  |  |
| Plasmid: pRDH10, *ori*(R6K) *mobRP4* *sacRB* Cm<sup>R</sup> Tet<sup>R</sup> | Kingsley et al., 1999 | pRDH10 |
| Plasmid: Up- and downstream region of *tar* (IR715) in pRDH10 | This study | pWJ39 |
| Plasmid: Up- and downstream region of *aspA* (IR715) in pRDH10 | This study | pWJ40 |
| Plasmid: Up- and downstream region of *dcuA* (IR715) in pRDH10 | This study | pWJ41 |
| Plasmid: Up- and downstream region of *frdABCD* (IR715) in pRDH10 | This study | pWJ51 |
| Plasmid: pKD46, Spec<sup>R</sup> P<sub>BAD</sub>-*gam-beta-exo oriR101 repA101<sup>ts</sup>* | Datsenko and Wanner, 2000 | pKD46 |
| Plasmid: pKD3, Carb<sup>R</sup> FRT Cm<sup>R</sup> FRT PS1 PS2 *oriR6K<sub>γ</sub>* | Datsenko and Wanner, 2000 | pKD3 |
| Plasmid: pKD13, Carb<sup>R</sup> FRT Km<sup>R</sup> FRT PS1 PS4 *oriR6K<sub>γ</sub>* | Datsenko and Wanner, 2000 | pKD13 |
| **Software and algorithms** |  |  |
| GraphPad Prism 8 | GrapPad Software | N/A |

## SUPPLEMENTARY MATERIAL

**Table S1.**
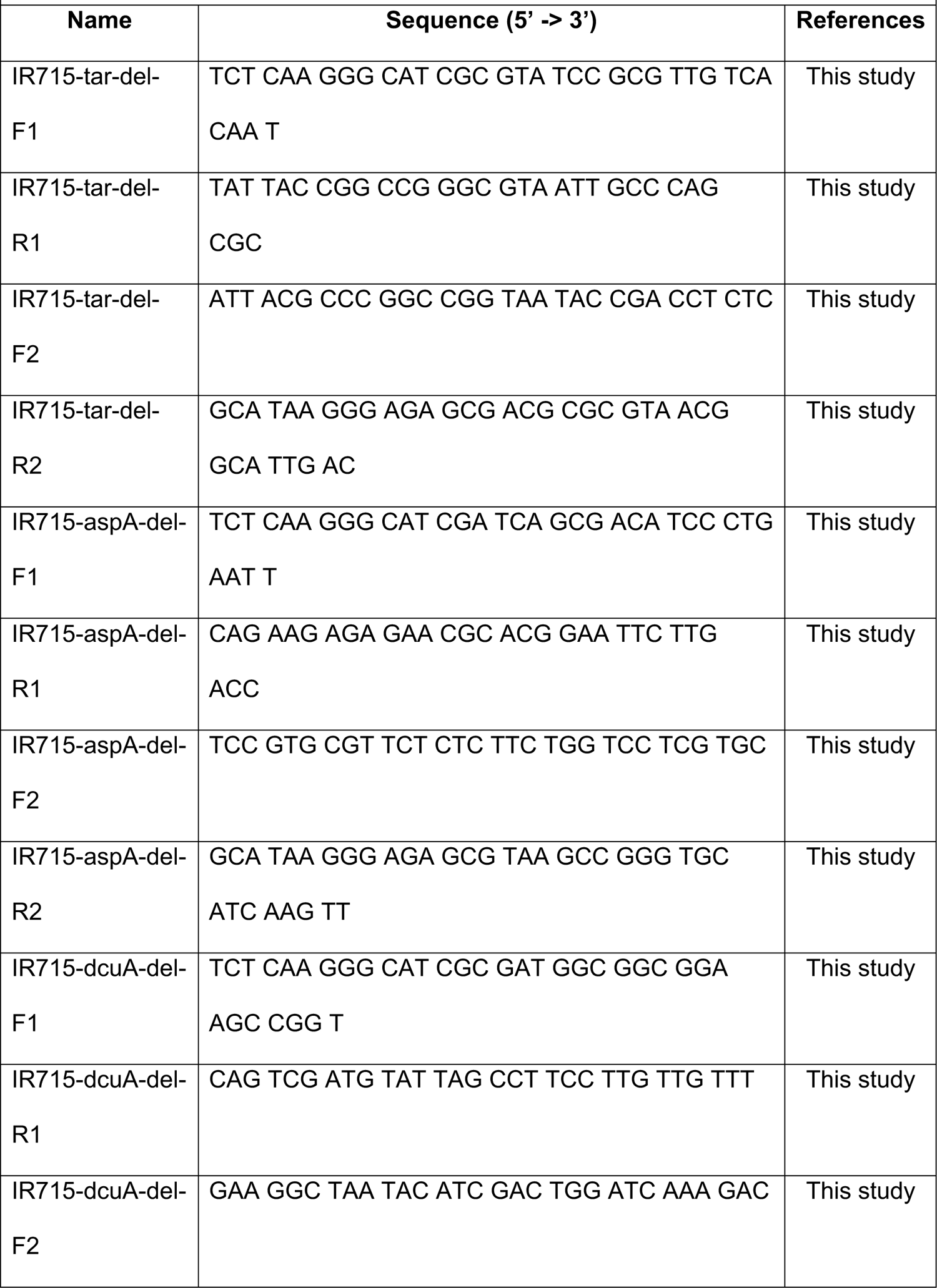
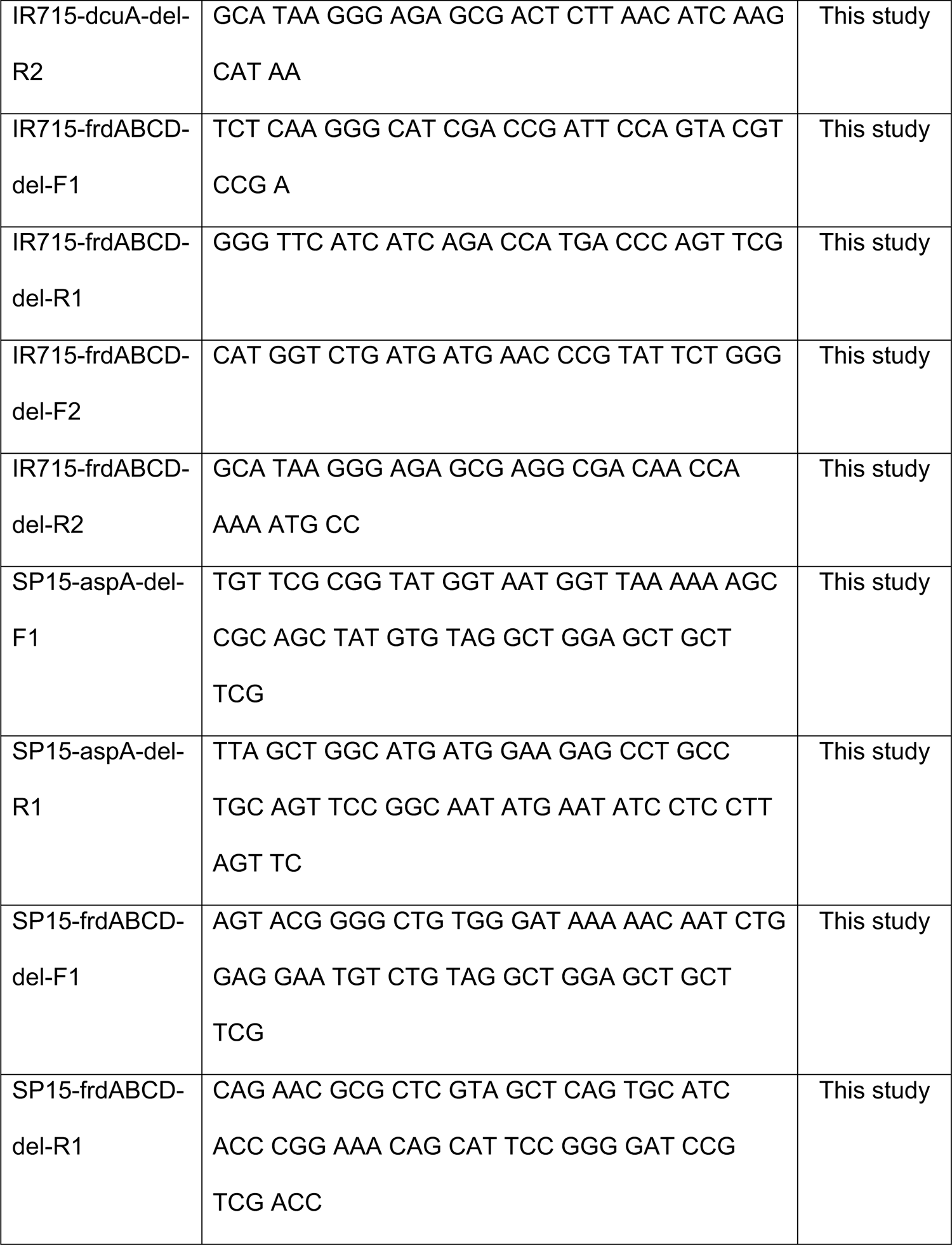
Primers used to construct plasmids and bacterial strains.

**Table S2.** Plasmids used to construct bacterial mutants.

| Table S2. Plasmids used to construct bacterial mutants. |  |  |
| --- | --- | --- |
| Name | Description | References |
| pRDH10 | *ori(R6K) mobRP4 sacRB Cm<sup>R</sup> Tet<sup>R</sup>* | Kingsley et al.,<br>1999 |
| pKD46 | Spec<sup>R</sup> P<sub>BAD</sub>-*gam-beta-exo oriR101 repA101<sup>ts</sup>* | Datsenko and<br>Wanner, 2000 |
| pKD3 | Carb<sup>R</sup> FRT Cm<sup>R</sup> FRT PS1 PS2 *oriR6K<sub>γ</sub>* | Datsenko and<br>Wanner, 2000 |
| pKD13 | Carb<sup>R</sup> FRT Km<sup>R</sup> FRT PS1 PS4 *oriR6K<sub>γ</sub>* | Datsenko and<br>Wanner, 2000 |
| pCP20 | Carb<sup>R</sup> Cm<sup>R</sup> *cl857 λP<sub>R</sub>flp oripSC101<sup>ts</sup>* | Datsenko and<br>Wanner, 2000 |
| pWJ39 | pRDH10::*Δtar* | This study |
| pWJ40 | pRDH10::*ΔaspA* | This study |
| pWJ41 | pRDH10::*ΔdcuA* | This study |
| pWJ51 | pRDH10::*ΔfrdABCD* | This study |

**Table S3.**
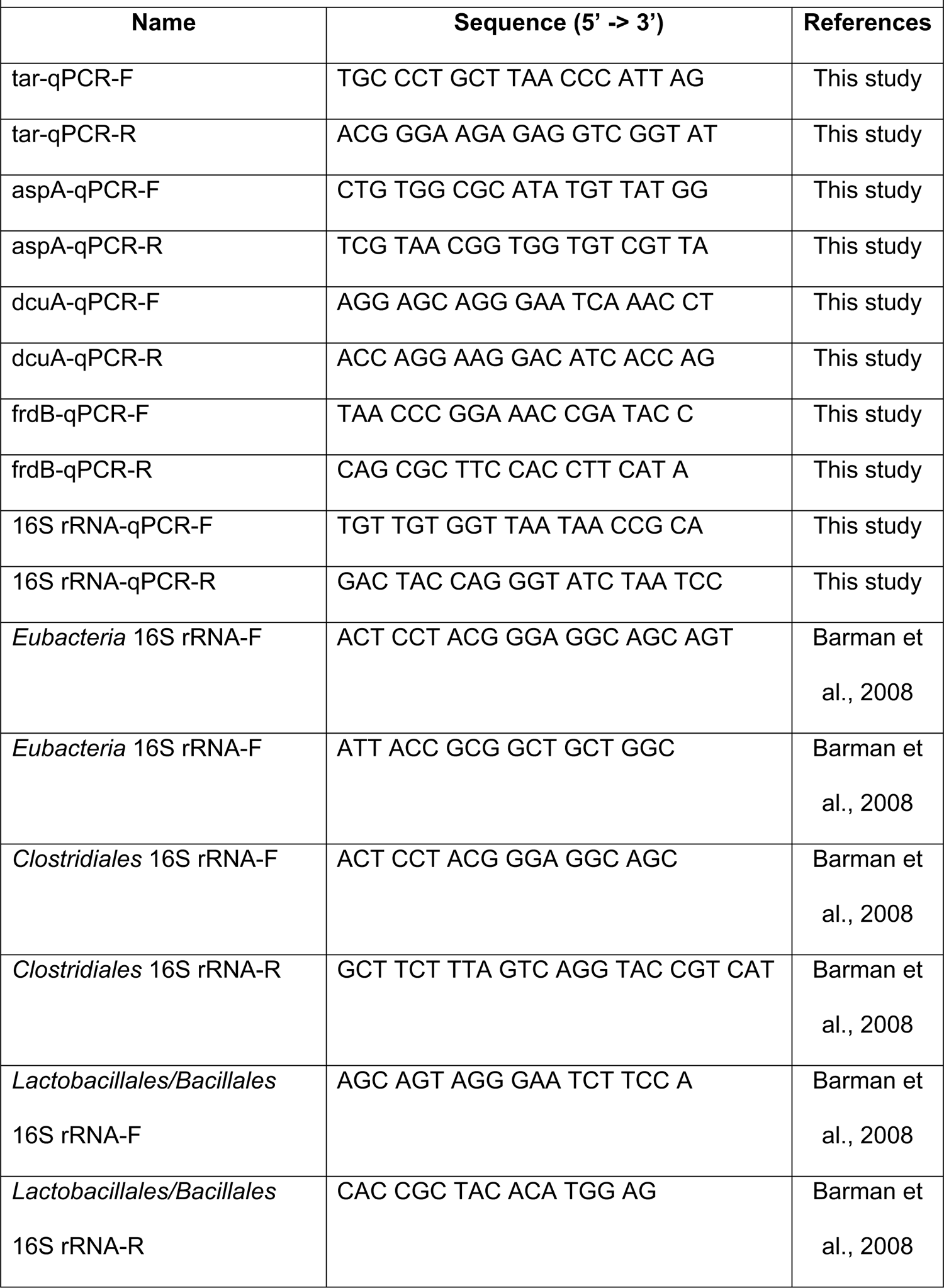
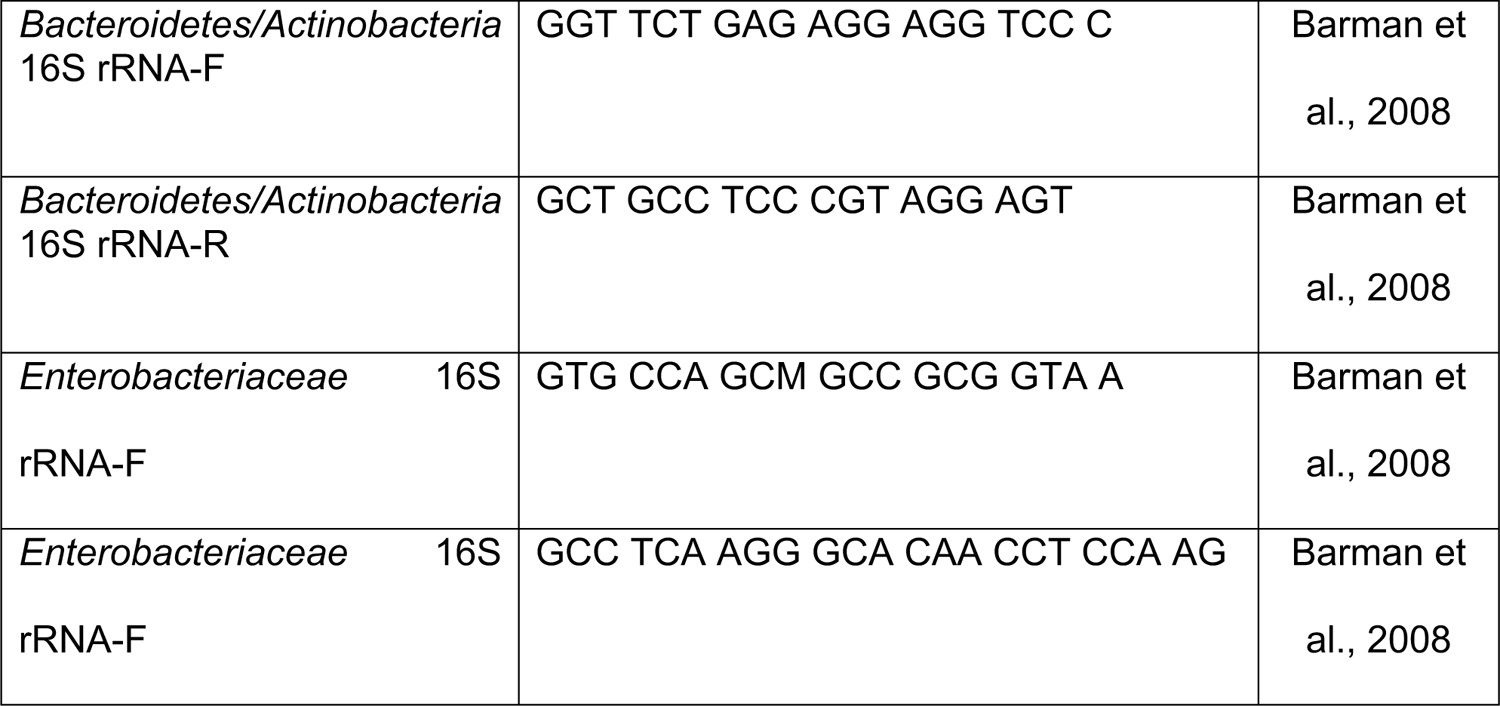
qRT-PCR primers for *Salmonella* gene expression.

**Table S4.**
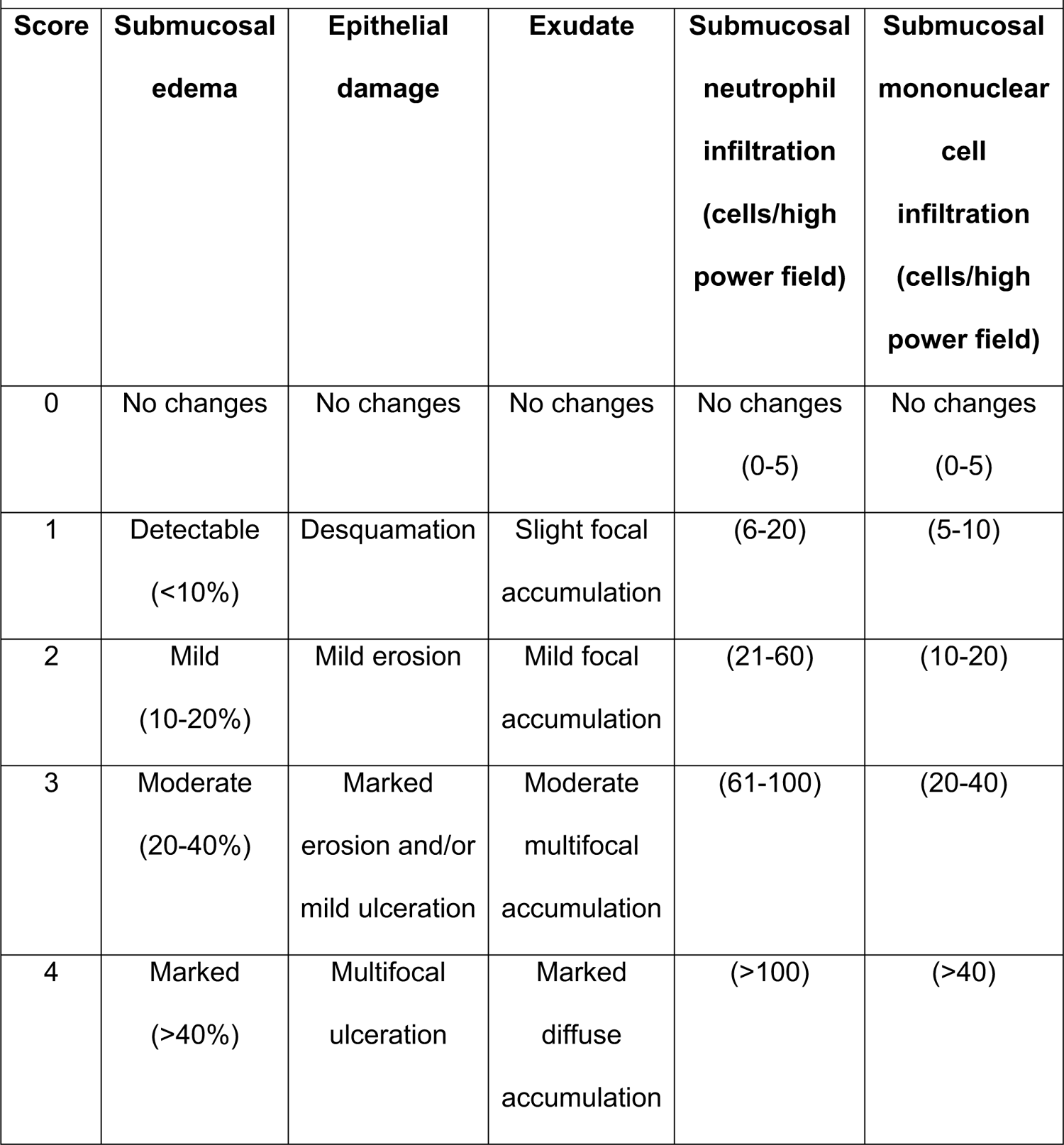
Criteria for blinded scoring of histopathological changes.

**Figure S1.**
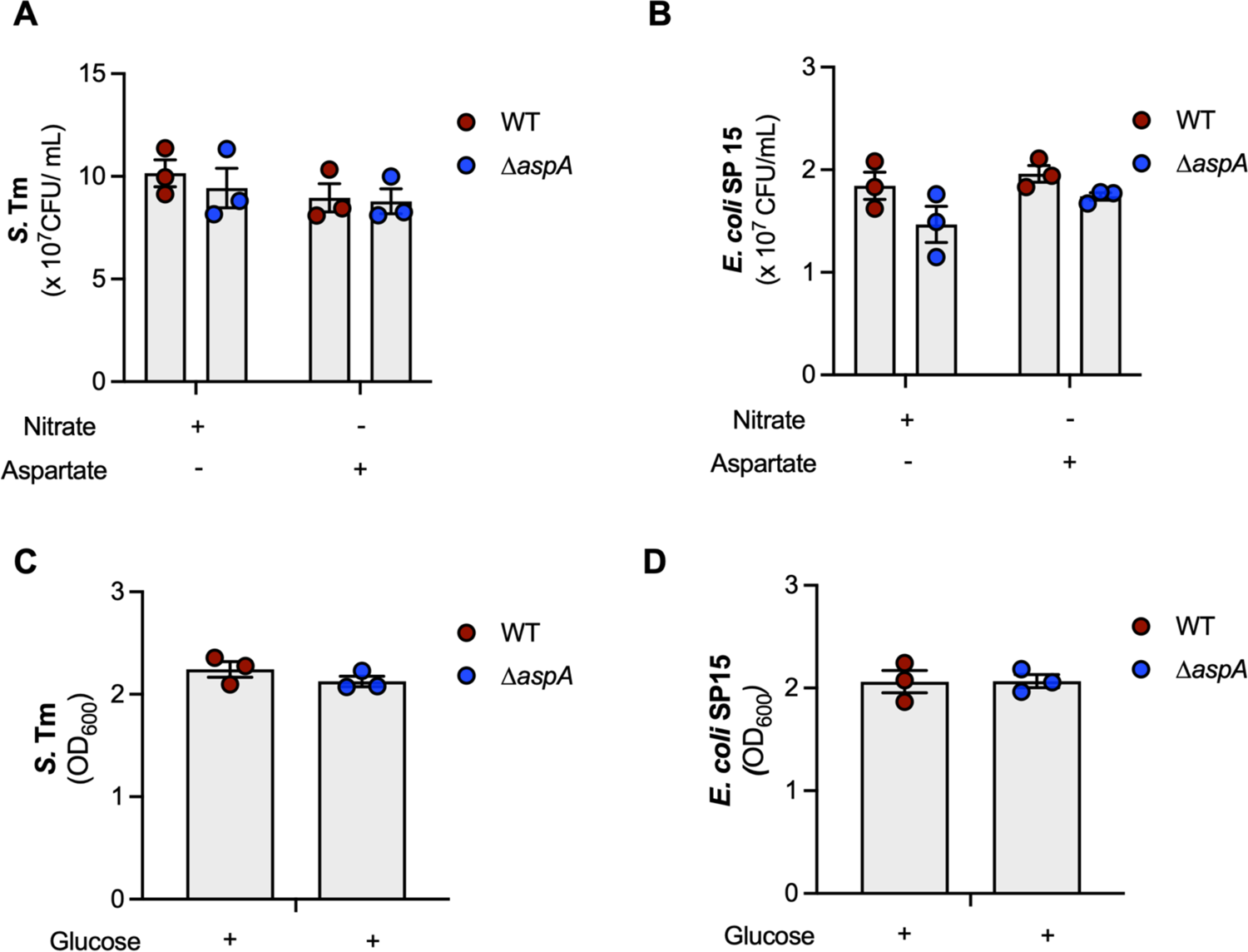
*Salmonella enterica* serovar Typhimurium and *Escherichia coli* SP15 Δ*aspA* mutants are not defective for growth on nitrate and glucose. Related to Figure 1. (A-B) Wild-type *S.* Tm (A) and *Escherichia coli* SP15 (B) and isogenic Δ*aspA* mutants were grown in NCE minimal media supplemented or not with L-aspartate (30mM) or sodium nitrate (40mM) for 18 hours under anaerobic conditions. (C-D) Wild-type *S.* Tm (A) and *Escherichia coli* SP15 (B) and isogenic *ΔaspA* mutants were inoculated with OD_600_ = 0.1 and then grown in NCE minimal media supplemented with glucose (5 mM) for 18 hours under anaerobic conditions. Bacterial growth was determined by measuring OD_600_. Each dot represents one biological replicate (average of triplicate technical replicate per biological replicate). Bars represent mean ± SEM.

**Figure S2.**
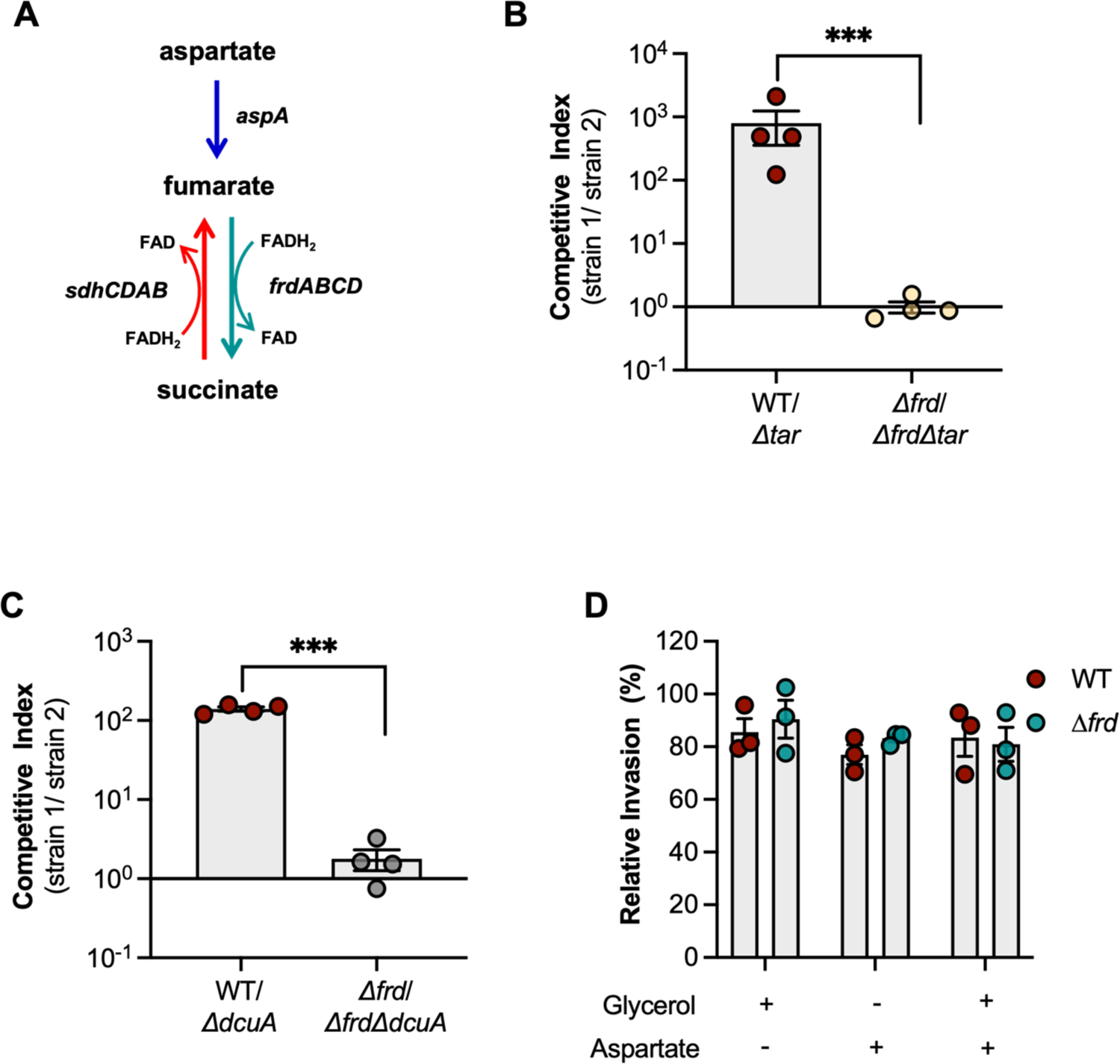
Chemotaxis towards exogenous aspartate and transport of the amino acid support fumarate-dependent anaerobic respiration *in vivo*. Related to Figure 3. (A) Schematics of aspartate-dependent fumarate respiration. (B) Female CBA/J mice were intragastrically infected with 10^9^ CFU of a 1:1 ratio of wild-type *S.* Tm (WT) and isogenic Δ*tar* mutant or a 1:1 ratio of *S.* Tm *Δfrd* and isogenic *Δfrd/Δtar* mutant for ten days. Competitive index in fecal samples was determined at day ten post-infection. (C) Female CBA/J mice were intragastrically infected with 10^9^ CFU of a 1:1 ratio of wild-type *S.* Tm (WT) and isogenic *ΔdcuA* mutant or a 1:1 ratio of *S.* Tm *Δfrd* and isogenic *Δfrd/ΔdcuA* mutant for ten days. Competitive index in fecal samples was determined at day ten post-infection. (D) Invasion essay in Caco-2 cells of wild-type *S.* Tm (WT) and isogenic *Δfrd* strain. MOI=10. For *in vitro* experiments, each dot represents one biological replicate (average of triplicate technical replicate per biological replicate). For mouse experiments, each dot represents data from one animal (biological replicate). Bars represent mean ± SEM. ***, p < 0.001.

**Figure S3.**
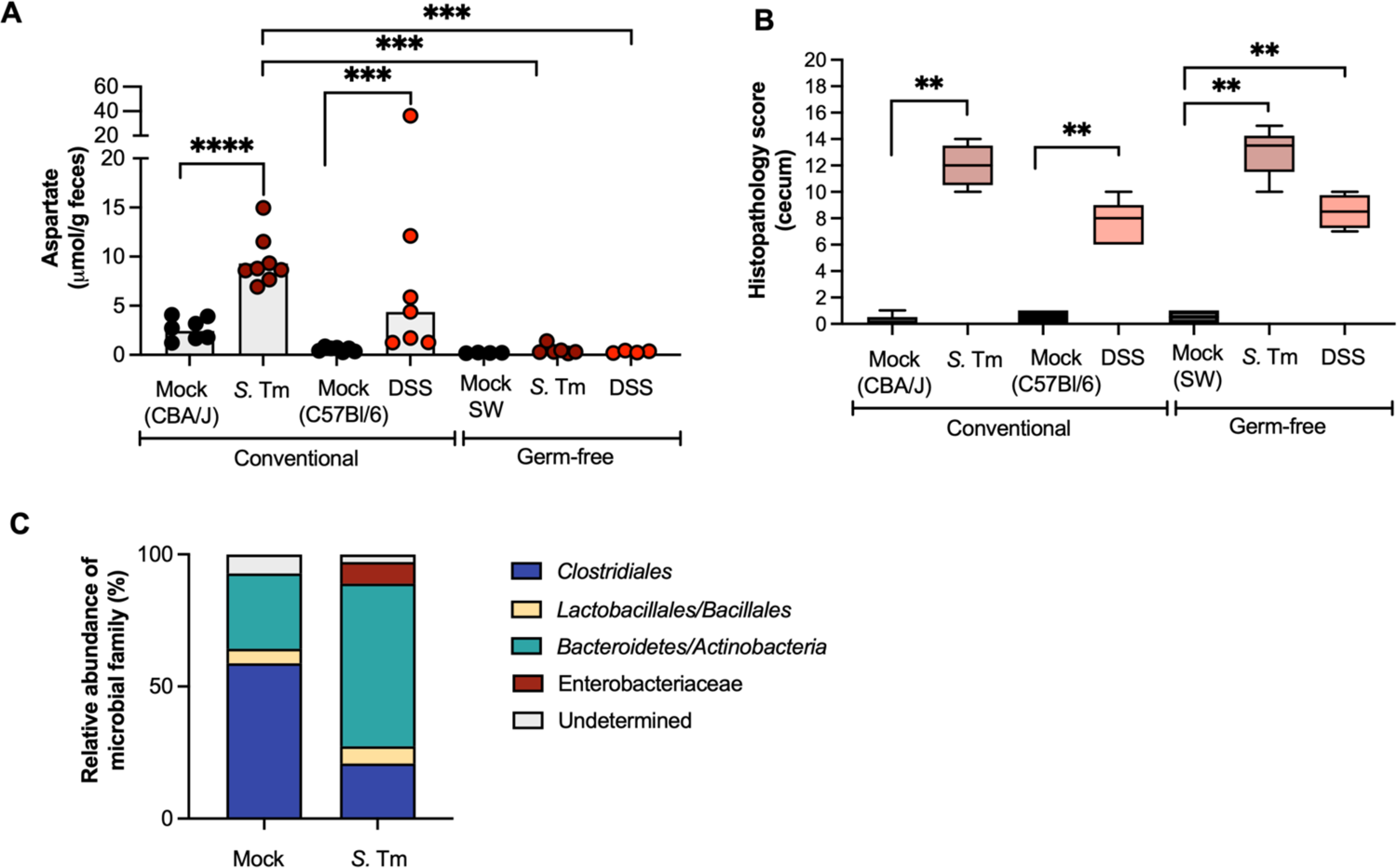
Colitis does not result in increased levels of aspartate in the intestinal lumen of germ-free mice. Related to Figure 5. (A) Conventional CBA/J and germ-free Swiss Webster (SW) mice were intragastrically infected with 10^9^ CFU of wild-type *S.* Tm. Conventional C57Bl/6J mice and germ-free SW mice were given 2.5% DSS in the drinking water for seven days to induce colitis. Fecal aspartate levels were measured at day ten post-infection for conventional CBA/J mice, at day three post-infection for germ-free SW mice and at day eight after beginning of treatment for DSS-treated mice. (B) Combined histopathology score of pathological lesions in the cecum of mice from (A) at day ten post-infection for conventional CBA/J mice, at day three post-infection for germ-free SW mice and at day eight after beginning of treatment for DSS-treated mice. (n=5-8). (C) Relative abundance of microbiota members performed by real-time qPCR in fecal samples of female CBA/J mice infected with 10^9^ CFU of wild-type *S.* Tm for ten days. Each dot represents data from one animal (biological replicate). Bars represent mean ± SEM. The boxes in the whisker plot represent the first to third quartiles, and the mean value of the gross pathology scores is indicated by a line. **, p < 0.01; ***, p < 0.001.

**Figure S4.**
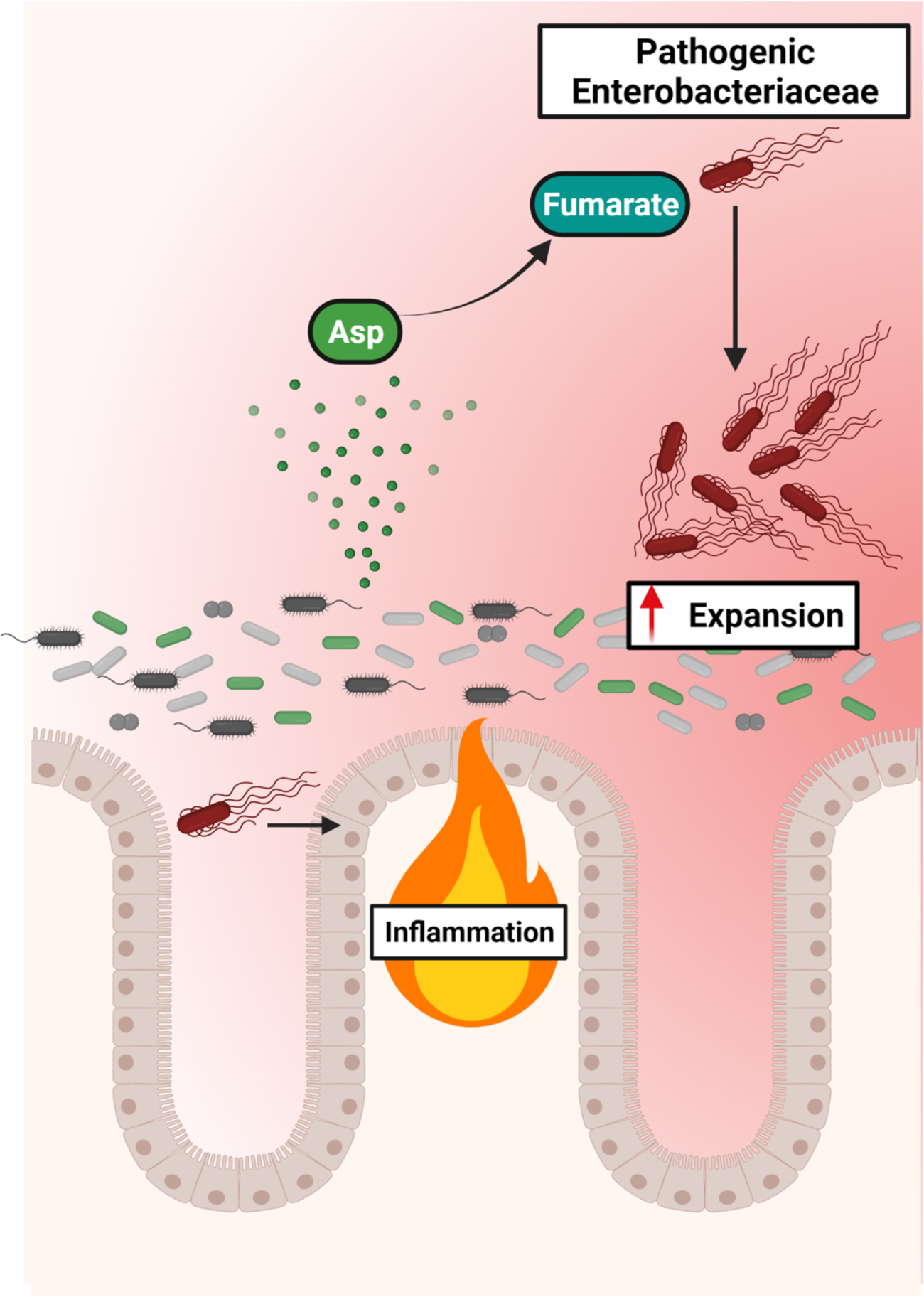
Pathogenic Enterobacteriaceae rely on microbiota-derived aspartate to support fumarate-dependent fumarate respiration in the inflamed gut. Created with BioRender.com.

